# TNF deficiency dysregulates inflammatory cytokine production leading to lung pathology and death during respiratory poxvirus infection

**DOI:** 10.1101/2019.12.22.883728

**Authors:** Ma. Junaliah Tuazon Kels, Esther Ng, Zahrah Al Rumaih, Pratikshya Pandey, Sigrid R. Ruuls, Heinrich Korner, Timothy P. Newsome, Geeta Chaudhri, Gunasegaran Karupiah

## Abstract

Excessive tumor necrosis factor (TNF) is known to cause significant pathology. Paradoxically, deficiency in TNF (TNF^-/-^) also caused significant pathology during respiratory ectromelia virus (ECTV) infection, a surrogate mouse model for smallpox. TNF^-/-^ mice succumbed to fulminant disease whereas wild-type mice, and those expressing only transmembrane TNF, recovered. TNF deficiency did not affect viral load or leukocyte recruitment but caused severe lung pathology and excessive production of the cytokines IL-6, IL-10, TGF-β, and IFN-γ. Blockade of these cytokines reduced lung pathology concomitant with induction of protein inhibitor of activated STAT3 (PIAS3) and/or suppressor of cytokine signaling 3 (SOCS3), factors that inhibit STAT3 activation. Short-term inhibition of STAT3 activation in ECTV-infected TNF^-/-^ mice with an inhibitor reduced lung pathology. TNF is essential for regulating inflammation and its deficiency exacerbates ECTV infection as a consequence of significant lung pathology caused by dysregulation of inflammatory cytokine production, in part via overactivation of STAT3.

## INTRODUCTION

Tumor necrosis factor (TNF) plays essential roles in normal physiology (Uysal et al., 1997), maintenance of immune homeostasis and acute inflammation (Arnett et al., 2001; Korner et al., 1997; Marino et al., 1997; Papathanasiou et al., 2015; Pasparakis et al., 1996), and anti-microbial defense (Bean et al., 1999; Domm et al., 2008; Kalliolias and Ivashkiv, 2016; Wilhelm et al., 2001). It has also been implicated in many of the detrimental effects of chronic inflammation in autoimmune conditions (Korner et al., 1995) and during infection (Belisle et al., 2010; Szretter et al., 2007; Tracey et al., 1987). The antimicrobial property of TNF has been underscored in patients receiving anti-TNF therapy to treat TNF-mediated inflammatory diseases, which has been shown to result in reactivation of some bacterial, parasitic, fungal and a limited number of viral infections (Domm et al., 2008; Garcia-Gonzalez et al., 2012; Kalliolias and Ivashkiv, 2016; Saunders and Britton, 2007; Subramaniam et al., 2013).

TNF is produced following activation of the nuclear factor-kappa B (NF-κB) inflammatory pathway, often in response to stress or infection. It is expressed as a transmembrane protein (mTNF), which is then cleaved by metalloproteinase enzymes to produce the secreted form, sTNF (Black et al., 1997; Moss et al., 1997). Both forms of TNF can mediate effector functions, but the relative contribution of each during viral infection is not known. TNF is known to contribute to an exaggerated immune response leading to host tissue destruction and immunopathology during some viral infections (Damjanovic et al., 2011; Peper and Van Campen, 1995).

Some members of the orthopoxvirus (OPV) family, like variola virus (agent of smallpox), monkeypox virus, cowpox virus and ectromelia virus (ECTV) encode several host response modifiers including tumor necrosis factor receptor (TNFR) homologs, suggesting an important role for TNF in protective immunity (Alcami, 2003; Alejo et al., 2009; Herbein and O’Brien, 2000; Loparev et al., 1998; Rahman and McFadden, 2006; Seet et al., 2003). ECTV is a mouse pathogen that causes mousepox, a disease very similar to smallpox, and one of the best small-animal models available for investigating the pathogenesis of OPV infections. Indeed, there are at least two lines of evidence indicating that TNF plays a crucial role in protection and recovery of mice from OPV infections. First, TNF production is strongly associated with resistance to ECTV infection in mice (Chaudhri et al., 2004). Resistant mice like the wild-type (WT) C57BL/6 strain produce high levels of TNF, which is associated with potent inflammatory and immune responses whereas the susceptible BALB/c strain produces little TNF, associated with weak inflammatory and immune responses. Second, the ECTV encoded viral TNFR (vTNFR) homolog, termed cytokine response modifier D (CrmD), is known to modulate the host cytokine response (Alejo et al., 2018). Infection of the ECTV-susceptible BALB/c strain with a CrmD deletion mutant virus augmented inflammation, natural killer (NK) cell and cytotoxic T lymphocyte (CTL) activities, resulting in effective control of virus replication and survival from an otherwise lethal infection.

Excessive TNF production, via NF-κB activation, can induce the production of other pro-inflammatory cytokines, including interleukin 6 (IL-6), which in turn can activate signal transducer and activator of transcription 3 (STAT3) (Zhong et al., 1994). Both NF-κB and STAT3 signaling pathways are closely intertwined and regulate an overlapping group of target genes including those associated with inflammation (Goldstein et al., 2017; Kasembeli et al., 2018; Ma et al., 2017; Oeckinghaus et al., 2011). STAT3 activation occurs through phosphorylation to form phosphorylated STAT3 (pSTAT3), and its activity is regulated through dephosphorylation by phosphatases (Hillmer et al., 2016; O’Shea et al., 2015) or by inhibitors of STAT3, namely protein inhibitor of activated STAT3 (PIAS3) (Chung et al., 1997; Shuai and Liu, 2005; Yagil et al., 2010) and suppressor of cytokine signaling 3 (SOCS3) (Babon et al., 2014; Carow and Rottenberg, 2014).

Using the mousepox model, we show here that TNF has no antiviral effects in C57BL/6 mice but plays a key role in regulating inflammatory cytokine production and resolution of lung inflammation during a respiratory infection. C57BL/6 WT mice and mice expressing only the non-cleavable transmembrane form of TNF (mTNF^Δ/Δ^) recovered from ECTV infection, whereas TNF-deficient (TNF^-/-^) mice succumbed with uniform mortality. TNF deficiency resulted in dysregulated production of IL-6, IL-10, transforming growth factor beta (TGF-β), and interferon gamma (IFN-γ) and caused significant lung pathology in virus-infected mice. Cytokine blockade with monoclonal antibodies (mAb) significantly reduced lung pathology contemporaneous with increased levels of PIAS3 and/or SOCS3 expression, suggesting that excessive cytokine production in TNF^-/-^ mice might be due to dysregulated STAT3 activation. Indeed, short-term treatment of ECTV-infected TNF^-/-^ mice with mAb against IL-6, IL-10 receptor (IL-10R), TGF-β, or IFN-γ or an inhibitor of STAT3 activation significantly reduced lung pathology. Long-term treatment with anti-IL-6 or anti-TGF-β but not anti-IL-10R or STAT3 inhibitor resulted in the recovery of mice and effective virus control. Our findings indicate that during respiratory OPV infection, TNF deficiency results in dysregulated cytokine production, in part due to overactivation of STAT3 signaling, causing massive lung pathology and death.

## RESULTS

### mTNF is induced rapidly in WT mice and TNF deficiency exacerbates respiratory ECTV infection independent of viral load or cell-mediated immunity

In WT mice infected with ECTV through the intranasal (i.n.) route, TNF mRNA was detectable in lungs at day 3 post-infection (p.i.), with levels still relatively high at day 12 (Figure S1A). In lung homogenates of WT mice, sTNF was detectable from about day 7 p.i. (Figure S1B) whereas mTNF was detectable earlier but levels of both forms were highest at day 12 p.i. (Figure S1C).

WT, TNF^-/-^ and mTNF^Δ/Δ^ mice were next infected i.n. with 25 plaque forming units (PFU) of ECTV and assessed for clinical scores, weight loss, survival, viral load, NK cell and virus-specific CTL responses.

Evaluation of the clinical presentation using a scoring system (described in Table S1) indicated that TNF^-/-^ animals fared significantly worse than WT or mTNF^Δ/Δ^ mice beginning at day 8 p.i. (Figure 1A). Mild clinical manifestations of infection were evident in all three strains beginning at days 5 and 6 p.i., but only TNF^-/-^ mice showed postural changes and inactivity from day 7. All 3 strains exhibited piloerection, lacrimation, and nasal discharge from day 8 p.i., however, only TNF^-/-^ mice started to succumb to the infection, some with apparent dyspnea. All strains lost weight from day 7 p.i., however, WT and mTNF^Δ/Δ^ animals started to recover from day 12 p.i. (Figure 1B). One mTNF^Δ/Δ^ mouse died at day 9 whereas all TNF^-/-^ mice succumbed between days 9 and 11 p.i. (Figure 1C). In some experiments, TNF^-/-^ mice survived beyond 12 days but eventually succumbed to disease whereas WT and mTNF^Δ/Δ^ mice fully recovered.

**Figure 1.**
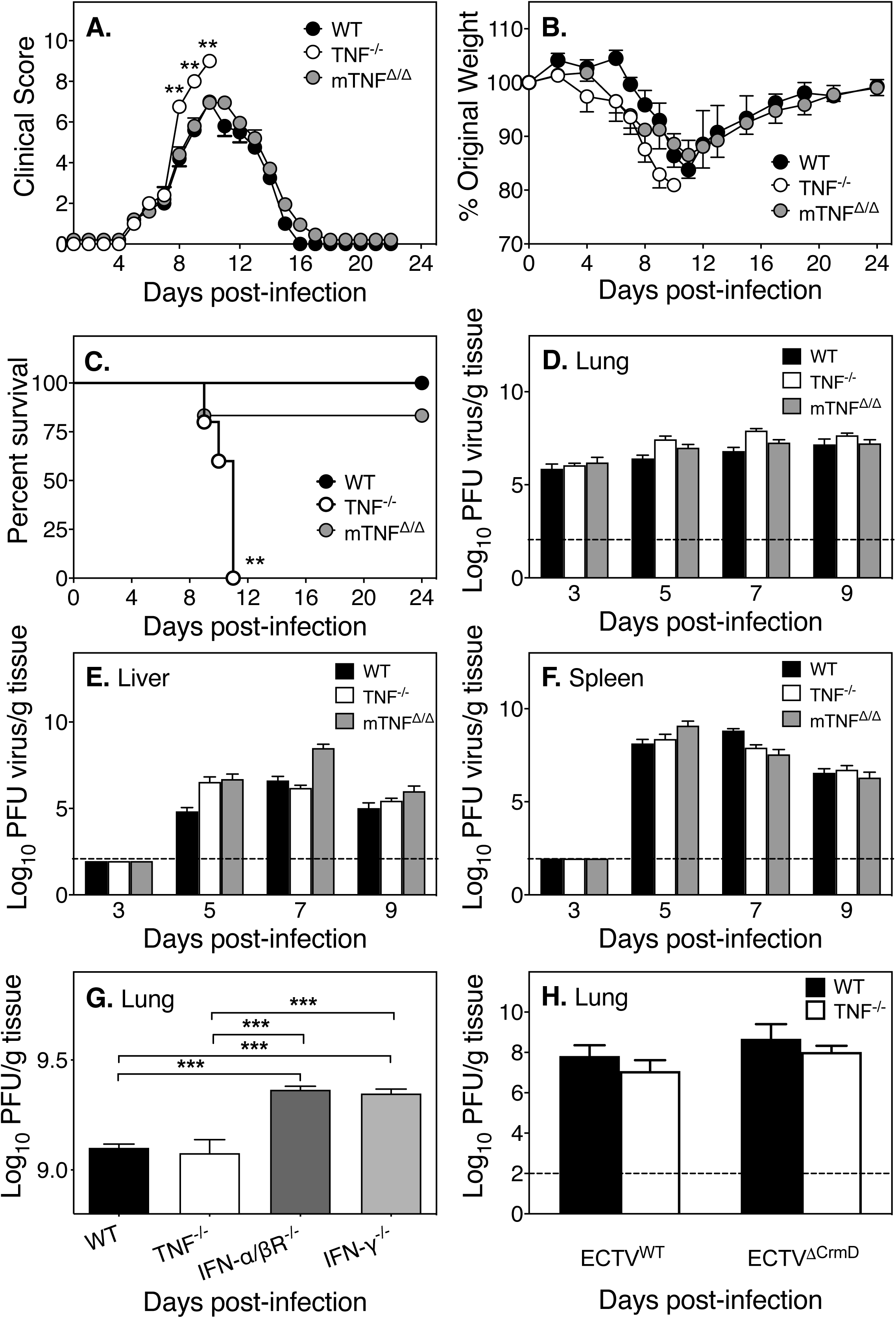
TNF is critical for recovery of mice from ECTV infection but is not required for control of virus replication. Groups of WT, TNF^-/-^ and mTNF^Δ/Δ^ mice were infected i.n. with 25 PFU ECTV. (A) Clinical scores (criteria detailed in Table S1), (B) weights and (C) survival were monitored during the course of infection. Additional groups of mice were infected with ECTV and 5 mice from each strain were sacrificed on the indicated days p.i. and viral load in (D) lungs, (E) livers, and (F) spleens were determined by plaque assay. (G) Lung viral load in groups of 5 WT, TNF^-/-^, IFN-α/βR^-/-^ and IFN-γ^-/-^ ECTV-infected mice at day 5 p.i. determined by plaque assay. (H) Lung viral load in groups of 5 WT and 5 TNF^-/-^ mice infected with either ECTV^WT^or ECTV^ΔCrmD^ at day 7 p.i. Data are expressed as means ± SEM. Statistical analysis was undertaken as follows: for (A), clinical scores for TNF^-/-^ and mTNF^Δ/Δ^ mice on each day p.i. were compared to WT mice on that day using two-tailed Mann-Whitney test; for (B) weights for TNF^-/-^ and mTNF^Δ/Δ^ mice on each day p.i. were compared to WT mice using multiple unpaired t-tests and the Holm-Sidak’s correction for multiple comparisons; for (C) survival curves were compared using Log-rank (Mantel-Cox) statistical test; and for (D)-(H), log transformed viral loads were analyzed by two-way ANOVA with Holm-Sidak’s correction for multiple comparison and the broken line corresponds to the limit of detection in plaque assays. *, p ≤ 0.05; **, p ≤ 0.01; ***, p ≤ 0.001; ****, p ≤ 0.0001. Results shown are representative of 3 independent experiments for (A)-(F), 1 experiment for (G) and 2 experiments for (H). See also Table S1.

Susceptibility to ECTV infection has been associated with uncontrolled virus replication in all strains of mice studied to date (Chaudhri et al., 2004; Karupiah et al., 1993). Although TNF^-/-^ mice were susceptible to ECTV, they did not have increased viral load in the lung (Figure 1D), liver (Figure 1E) or spleen (Figure 1F) compared to titers in either WT or mTNF^Δ/Δ^ animals at days 3, 5, 7 or 9 p.i. Conversely, mice deficient in IFN-α/β receptor (IFN-α/βR^-/-^) or IFN-γ (IFN-γ^-/-^), strains known to rapidly succumb to mousepox due to uncontrolled virus replication (Panchanathan et al., 2005), had significantly higher viral load in lungs compared to WT or TNF^-/-^ mice (Figure 1G). We investigated whether CrmD, the vTNFR homolog encoded by ECTV, affected virus replication using the Naval strain of WT (ECTV^WT^) and CrmD mutant (ECTV^ΔCrmD^) viruses (Alejo et al., 2018). WT and TNF^-/-^ mice infected with either of the viruses had comparable viral loads in lungs, indicating that vTNFR did not affect viral replication (Figure 1H).

Since early activation of NK cells and CTL responses are critical for virus control and recovery from mousepox (Chaudhri et al., 2004; Delano and Brownstein, 1995; Karupiah et al., 1996; Parker et al., 2007), we investigated the effect of TNF deficiency on these responses. NK cell responses in the lungs (Figures S2A and S2B) and spleens (Figures S2C and S2D) were similar in WT, mTNF^Δ/Δ^, and TNF^-/-^ strains. Likewise, TNF deficiency did not affect the antiviral CTL responses in the lungs (Figure S2E) or spleen (Figure S2F). Thus, unlike IFN-γ and IFN-α/β, TNF neither possesses antiviral activity nor influences NK cell or CTL responses in C57BL/6 mice infected with ECTV (Chaudhri et al., 2004).

To exclude the possibility that the increased susceptibility of TNF^-/-^ mice might be related to the dysregulation of organogenesis and spatial organization of lymphoid tissue known to occur during development (Korner et al., 1997; Neumann et al., 1996; Pasparakis et al., 1996; Pasparakis et al., 1997), we used neutralizing anti-TNF mAb to treat ECTV-infected WT mice. Compared to the control mAb-treated animals, mice treated with anti-TNF mAb exhibited significantly higher clinical scores (Figure S3A) and 50% of the animals succumbed by day 9 p.i. (Figure S3B). The histopathological scores were also higher in anti-TNF mAb-treated animals (Figure S3C) but there were no differences in viral load between the groups (Figure S3D). Representative lung histology sections from naïve animals and from ECTV-infected mice treated with either control or anti-TNF mAb are shown in Figure S3E. These results are consistent with those in TNF^-/-^ mice and further confirmed that TNF is critical for recovery of mice from ECTV infection but that the cytokine has no antiviral effects.

### Exacerbation of lung pathology in the absence of of TNF

An examination of lung histological sections indicated that there were minimal changes in the lungs of WT and mTNF^Δ/Δ^ mice on days 3, 5, and 7 p.i., whereas edema was visible at day 7 in TNF^-/-^ animals (Figure S4A). By day 9 p.i., edema and leukocyte extravasation were seen in histological sections of all strains. Resolution of lung pathology was evident in WT and mTNF^Δ/Δ^ mice by day 11, but pathology worsened in TNF^-/-^ mice. As the clinical, virological, immunological, and histological responses to ECTV infection were similar in WT and mTNF^Δ/Δ^ mice (Figure 1, and Figures S2 and S4A), we undertook further analysis of the response to ECTV infection only in WT and TNF^-/-^ mice.

Examination of the gross anatomy of uninfected and virus-infected lungs indicated TNF^-/-^ lungs were more congested, boggier, physically larger, and contained more and bigger necrotic lesions compared to WT lungs (Figure 2A). The lung wet-to-dry weight ratio allowed assessment of fluid extravasation into the lung interstitial space, and our results indicated that TNF^-/-^ lungs had significantly higher levels of fluid on days 7, 9, and 11 p.i. (Figure 2B). Protein leakage is an indicator of increased vascular permeability, and TNF^-/-^ lungs had significantly higher levels of protein in bronchoalveolar lavage fluid (BALF) at day 11 p.i. compared to the WT lungs (Figure 2C).

**Figure 2.**
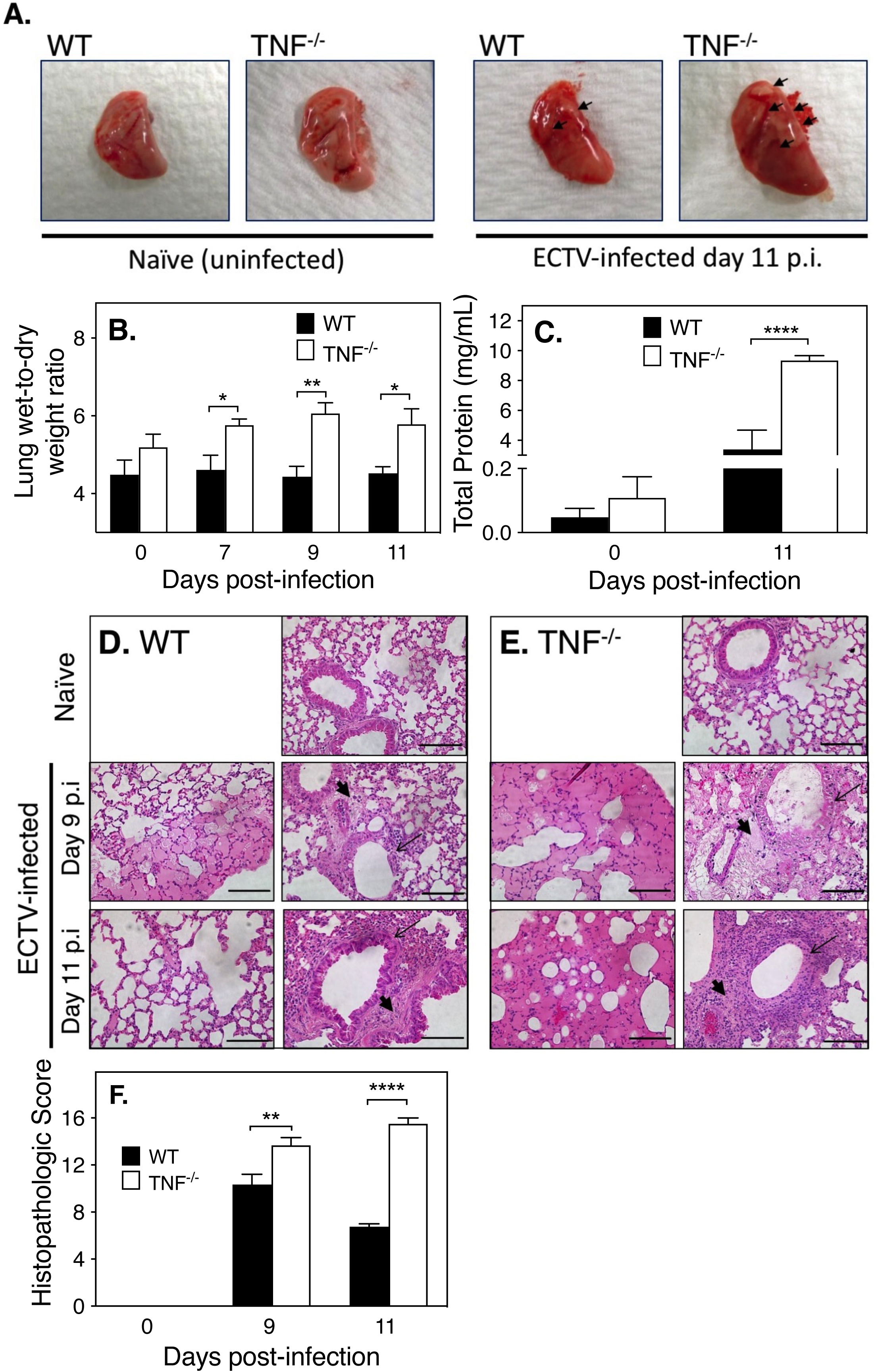
TNF deficiency in mice during ECTV infection exacerbates lung pathology. Groups of WT and TNF^-/-^ mice were infected with ECTV. Five mice from each group were sacrificed on days 7, 9, and 11 p.i, in addition to 5 uninfected mice of both genotypes. (A) Gross morphology of lungs from naïve WT and TNF^-/-^ mice is shown in comparison to lungs from day 11 p.i. Areas of consolidation and necrosis on the surface of the lungs are marked with arrows. (B) Fluid extravasation in the lungs was measured by the lung wet-to-dry weight ratios on the days indicated, and (C) protein concentration in BALF of naïve mice (day 0) and day 11 p.i. Histological changes in naïve and ECTV-infected (D) WT and (E) TNF^-/-^ mice at days 9 and 11 p.i. Slides were examined on all fields at 400x magnification and the bars shown correspond to 100 µm. Parenchymal edema, perivascular edema (thick arrows) and bronchial epithelial necrosis (thin arrows) are shown. (F) Lung histopathological scores on days 0, 9 and 11 p.i. using a scoring system described in Table S2. Data are expressed as means ± SEM. For (B), (C) and (F), statistical analysis was undertaken using two-way ANOVA followed by Holm-Sidak’s multiple comparisons test. Results are representative of 3 separate experiments. See also Figure S3 and Table S2.

Detailed microscopic examination of lung sections (Figures 2D and 2E) revealed that lung pathology was more severe in virus-infected TNF^-/-^ mice. There was intra-alveolar and perivascular edema in both strains but in the TNF^-/-^ group, the fluid extravasated to almost the entire lobe leading to alveolar septal wall collapse by day 9. By day 11 p.i., resolution of pathology was evident in WT lungs, wherein edema in the parenchyma started to clear, and restoration of the bronchial epithelial architecture was evident. In contrast, alveolar septal wall damage and edema in TNF^-/-^ lungs continued to increase from day 9 to day 11 p.i., parenchymal edema worsened, and the bronchial epithelia were completely denuded. The histopathological differences between the infected lungs of both mouse strains were semi-quantified using a scoring system (described in Table S2). TNF^-/-^ lungs had significantly higher scores than WT lungs on days 9 and 11 p.i. (Figure 2F). In WT mice, the histological scores were lower at day 11 compared to day 9, consistent with recovery of this strain from ECTV infection. TNF deficiency resulted in significant lung tissue damage and pathology, suggesting that TNF plays a role in anti-inflammatory processes.

### TNF is produced by many cell types and its deficiency does not affect leukocyte recruitment to the lungs but results in dysregulated inflammatory cytokine production

Cell types responsible for TNF production within the infected lungs were identified by immunohistochemistry (Figures 3A and 3B), and these were observed in the bronchial epithelium and alveoli where the virus antigen was also present (Figure S4B). Although morphologically identified leukocyte subsets were present in both strains of mice, as expected TNF protein was not detected in TNF^-/-^ lungs (Figure 3B). Lungs from WT mice showed that TNF protein was produced in response to infection by bronchial epithelial cells, and by multiple leukocyte subsets (Figure 3A). Flow cytometric analysis of digested lungs indicated that the total number of leukocytes (CD45^+^; Figure S5A) or individual leukocyte subsets (Figure 3C) were similar in both TNF^-/-^ and WT strains. Histological analysis of TNF^-/-^ lungs suggested that there was a confluence of leukocytes in specific areas at day 11 p.i. (Figure 2E), but the total number of leukocytes within the lungs had decreased compared to day 9 p.i. (Figures S5A-S5C). The consolidation of leukocytes in infected TNF^-/-^ mice was therefore not related to an increase in their numbers *per se* but may have been due to dysregulated cytokine production. We, therefore, measured levels of some specific inflammatory and regulatory cytokines in uninfected and virus-infected lung tissue.

**Figure 3.**
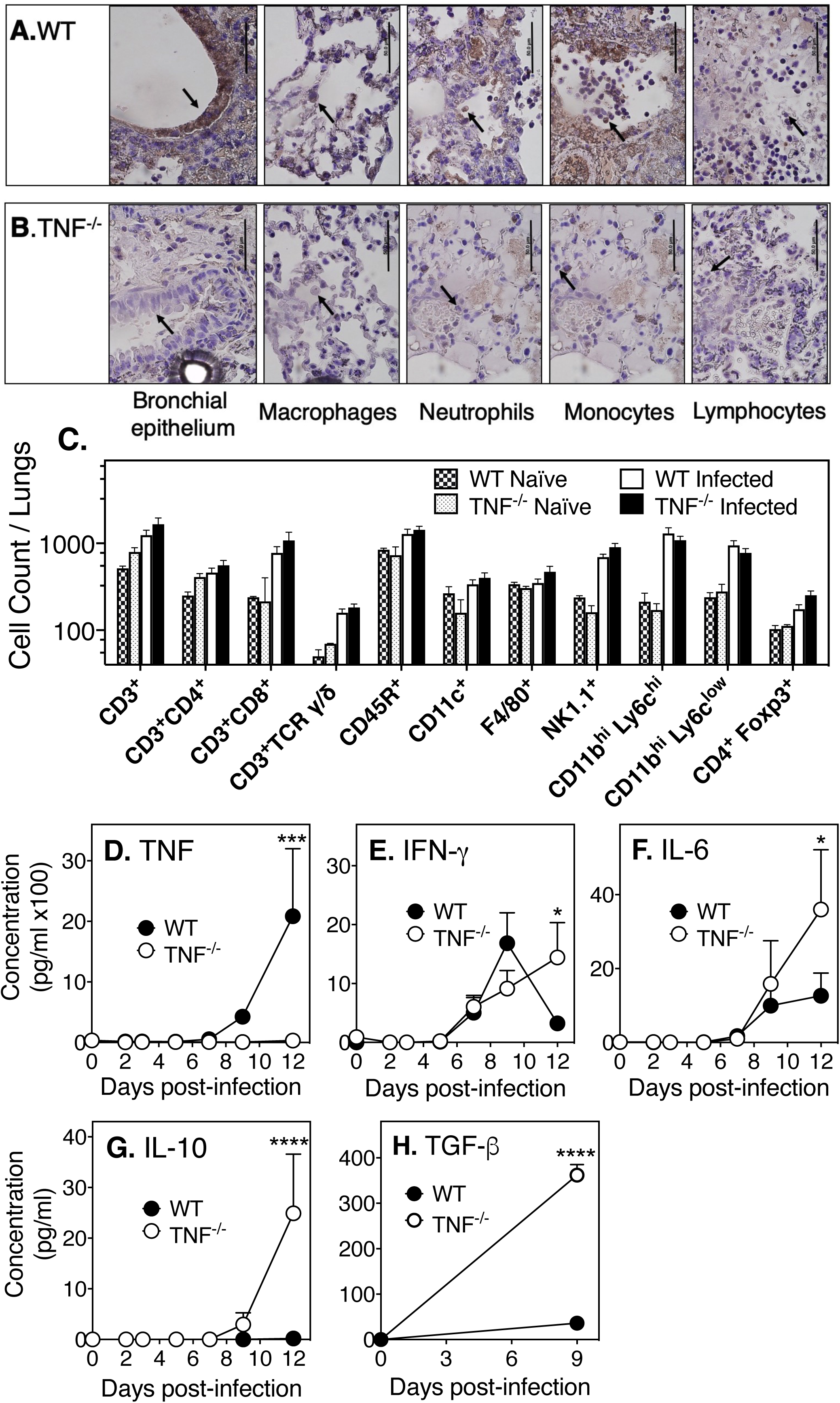
TNF deficiency does not affect lung leukocyte recruitment but results in increased production of specific inflammatory and regulatory cytokines. Immunohistochemistry of lung sections for TNF expression by bronchial epithelial cells and leukocyte subsets in ECTV-infected (A) WT and (B) TNF^-/-^ mice on day 9 p.i. Slides were examined at 1000x magnification. Arrows point to bronchial ephithelial cells and different cell types in the alveoli and bars correspond to 50 µm. (C) Flow cytometry analysis of digested lungs from naïve and ECTV-infected WT and TNF^-/-^ mice at day 9 p.i. Leukocyte subsets were identified by CD45.2 expression and one or more specific phenotypic markers as shown. Concentrations of (D) TNF, (E) IFN-γ, (F) IL-6, and (G) IL-10 in lung homogenates determined by Cytokine Bead Array. (H) TGF-β concentrations measured by ELISA. For (C)-(H), data are expressed as means ± SEM and were analyzed by two-way ANOVA followed by Holm-Sidak’s multiple comparisons test. Results are representative of 3 separate experiments with 5 animals per group. See also Figures S1B and S5 and Table S2. Data shown in panel 3D is the same as that shown in Figure S1B.

TNF protein was not detected in TNF^-/-^ animals whereas it was detectable in WT mice at day 9 p.i., and levels increased substantially by day 12 p.i. (Figure 3D). This was also reflected by TNF mRNA expression in WT animals (Figure S5D). Levels of IFN-γ, IL-6, IL-10, and TGF-β (Figures 3E-H) were significantly higher in lungs of TNF^-/-^ mice compared to WT animals at day 12 p.i. The higher levels of cytokine protein observed in the lungs of infected TNF^-/-^ mice were also reflected by increased mRNA levels of IFN-γ, IL-6, IL-10, and TGF-β (Figures S5E-H). In summary, ECTV infection of TNF^-/-^ mice resulted in increased production of a number of inflammatory and regulatory cytokines with the potential to cause lung damage.

### Blockade of specific cytokine function dampens lung pathology

To ascertain whether the increased production of cytokines identified in Figure 3E-H contributed to lung pathology, we reasoned that it might be possible to ameliorate the pathology by neutralizing or blockading the function of each cytokine with specific mAb *in vivo*. We used mAb against IFN-γ, IL-6, IL-10R or TGF-β to block cytokine function at day 7 p.i., co-incident with histological evidence of pulmonary edema (Figure S4A), and sacrificed mice at day 9. A single dose of mAb administered at this time also potentially minimized neutralization of cytokines known to play important antiviral roles early during the course of ECTV infection (Chaudhri et al., 2004; Ivashkiv, 2010; Karupiah et al., 1993; O’Gorman et al., 2010; Parker et al., 2007).

Treatment with control mAb (rat IgG) had minimal impact on disease outcome and, as expected, the lung pathology in WT mice (Figures 4A and 4B) was not as severe as in TNF^-/-^ mice (Figures 4K and 4L). Blockade of IFN-γ, IL-6, IL-10 or TGF-β function clearly reduced lung pathology and edema, including the amelioration of damage to the epithelial lining of bronchioles in both WT (Figures 4C-J) and TNF^-/-^ (Figures 4M-T) mice. Despite some improvement in lung pathology in mAb-treated WT mice, the histopathological scores were not significantly different from the control mAb-treated group (Figure 4U), possibly because WT mice did not produce high levels of these inflammatory cytokines (Figure 3) and the lung pathology was not as severe as in the TNF^-/-^ mice. Except for anti-IL-6 treatment, which reduced viral titers, mAb treatment did not affect viral load in WT mice (Figure 4V).

**Figure 4.**
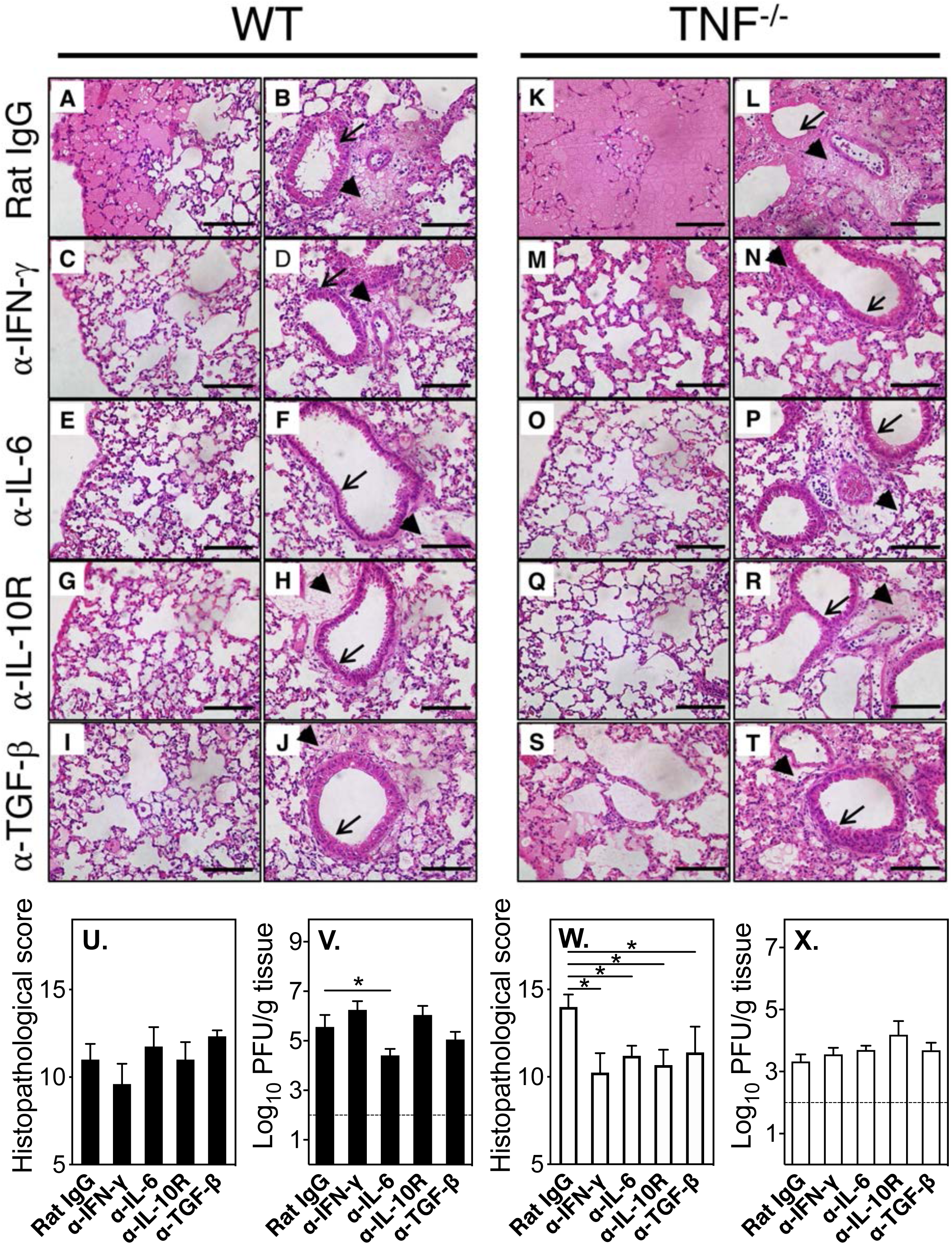
Blockade of TNF, IFN-γ, IL-6, IL-10R or TGF-β with mAb dampens lung pathology. Groups of WT (A-J) and TNF^-/-^ (K-T) ECTV-infected mice were treated on day 7 p.i. with isotype control rat IgG mAb (A, B, K, L), or specific mAb against IFN-γ (C, D, M, N), IL-6 (E, F, O, P), IL-10R (G, H, Q, R), or TGF-β (I, J, S, T) at 500 µg i.p. and sacrificed on day 9 p.i., lungs were collected, fixed, sectioned, H&E-stained and examined on all fields at 400x magnification. Perivascular edema (thick arrows) and bronchial epithelia (thin arrows) are shown. Bars correspond to 100 µm. In separate experiments, groups of WT (U-V) and TNF^-/-^ (W-X) mice were infected with ECTV and treated with mAb as above. Histopathological scores and viral load in WT mice (U and V) and TNF^-/-^ mice (W and X) at day 9 p.i. For panels V and X, the broken line correspond to the limit of virus detection. Data are expressed as means ± SEM and analyzed using unpaired t-test for (U) and (W). For (V) and (X), data were log transformed and analyzed by two-way ANOVA with Holm-Sidak’s correction for multiple comparison. Results are representative of 2 separate experiments with 5 animals per group.

In contrast, in TNF^-/-^ mice, treatment with mAb to IFN-γ, (Figures 4M and 4N), IL-6 (Figures 4O and 4P), IL-10R (Figures 4Q and 4R), or TGF-β (Figures 4S and 4T) significantly reduced histopathological scores (Figure 4W). The improved lung pathology in anti-cytokine mAb-treated mice was not associated with a reduction in viral titers (Figure 4X). Although not a consistent finding, ECTV titers in TNF^-/-^ mice in this experiment were lower than in the corresponding WT groups, further indicating that the worsened lung pathology in the mutant strain was not related to the viral load.

### *In vivo* neutralization of cytokine function increases pSTAT3, PIAS3, and SOCS3 levels in the lungs of infected mice

Neutralization of the function of four different cytokines, i.e. IFN-γ, IL-6, IL-10 or TGF-β led to the convergent result of dampening lung pathology (Figure 4), raising the possibility that their activities may regulate a common cytokine-signaling pathway. It is known that phosphorylation of STAT3 may be induced by cytokines like IL-6 (Zhong et al., 1994), TGF-β (McGeachy et al., 2007; Yamamoto et al., 2001), IL-10 (Niemand et al., 2003) and IFN-γ (Qing and Stark, 2004). We reasoned that cytokine blockade may have reduced STAT3 signaling by decreasing levels of pSTAT3 expression or through the induction of PIAS3 or SOCS3.

Western blot analysis revealed that levels of the inactive form of STAT3 were similar in uninfected or infected mice regardless of mouse strain or mAb treatment (Figure 5A). Minimal levels of pSTAT3 were detected in uninfected animals, and they increased by about 2-fold in WT mice and about 5-fold in TNF^-/-^ mice following infection (Figures 5A and 5B). In WT mice, pSTAT3 levels increased further by about 2-fold with anti-IL-6 or anti-TGF-β treatment, and higher than 3-fold following treatment with anti-IFN-γ or anti-IL-10R (Figures 5A and 5B). On the other hand, in TNF^-/-^ mice, the increases in pSTAT3 levels above control IgG-treated groups were marginal.

**Figure 5.**
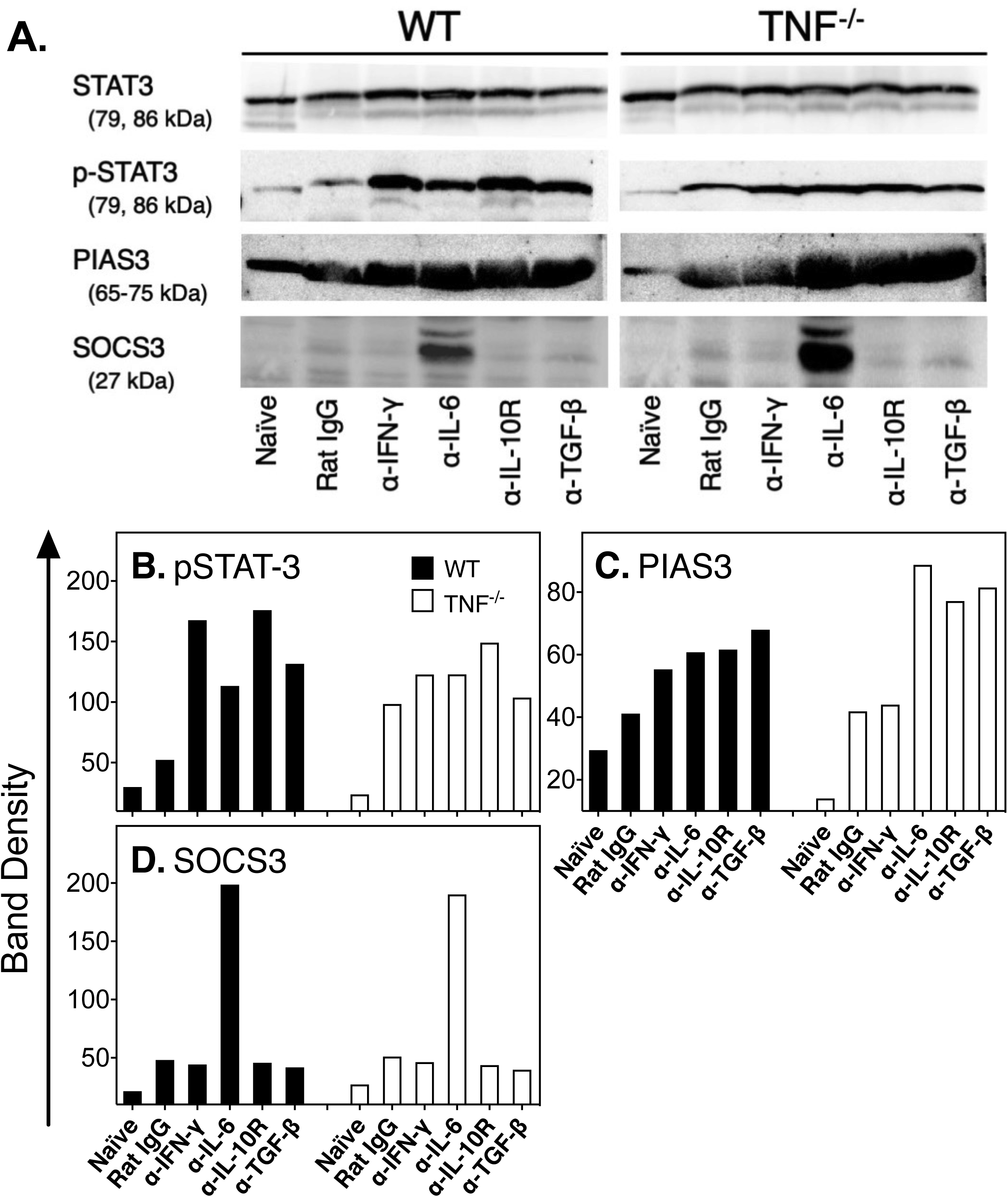
Short-term cytokine blockade increases levels of pSTAT3, PIAS3 and SOCS3 expression in ECTV-infected lungs. Groups of 5 WT and 5 TNF^-/-^ ECTV-infected mice were treated on day 7 p.i. with control rat IgG, or mAb against IFN-γ (α-IFN-γ), IL-6 (α-IL-6), IL-10R (α-IL-10R) or TGF-β (α-TGF-β) at 500 µg i.p. Lungs were collected on day 9 p.i. and homogenized for detection of STAT3, pSTAT3 (tyrosine-705 phosphorylation), PIAS3 and SOCS3 protein levels (A) by Western blot analysis. Band densities (B, C and D) were quantified using the ThermoScientific Pierce MYImageAnalysis Software. Analysis was done for individual mice and results shown are representative of one mouse per treatment group.

The basal levels of PIAS3 were about 3-fold lower in TNF^-/-^ mice compared with WT animals (Figures 5A and 5C). Infection augmented PIAS3 levels in both strains, with further increases noted after blockade of cytokine function (Figures 5A and 5C). In TNF^-/-^ mice, the PIAS3 band density increased by the largest magnitude following blockade of IL-6, IL-10R, and TGF-β. The increase in PIAS3 levels may have contributed to dampening of pSTAT3 activity and as a consequence, reduction in lung pathology.

Naïve animals did not express SOCS3, and this protein was detectable at only low levels after infection of both WT and TNF^-/-^ mice (Figures 5A and 5D). Only treatment with mAb against IL-6 resulted in significant increases in levels of SOCS3 (Figures 5A and 5D). It is likely that the combined actions of increased levels of PIAS3 and SOCS3 in the anti-IL-6-treated mice contributed to dampening of STAT3 activation and to effective amelioration of lung pathology and survival of mice.

### Inhibition of STAT3 activation ameliorates lung pathology

The preceding data suggested that it might be possible to dampen lung pathology through inhibition of STAT3 activation. Treatment with S3I-201, a selective STAT3 inhibitor that blocks STAT3 phosphorylation and dimerization (Chung et al., 1997), 7 days after ECTV infection resulted in a significant reduction in clinical scores in WT (Figure 6A) and TNF^-/-^ (Figure 6B) mice, and lung histopathological scores in both strains at day 9 p.i. (Figure 6C). However, the treatment did not have any effect on viral load in either strain (Figure 6D). Microscopically, STAT3 inhibitor treatment improved lung pathology in both WT (Figure 6E) and TNF^-/-^ (Figure 6F) lungs, and the effects were similar to blockade of specific cytokines (Figure 4). These results indicate that dysregulated inflammatory cytokine production and generation of lung immunopathology in the absence of TNF function is associated, at least in part, with hyperactivation of STAT3.

**Figure 6.**
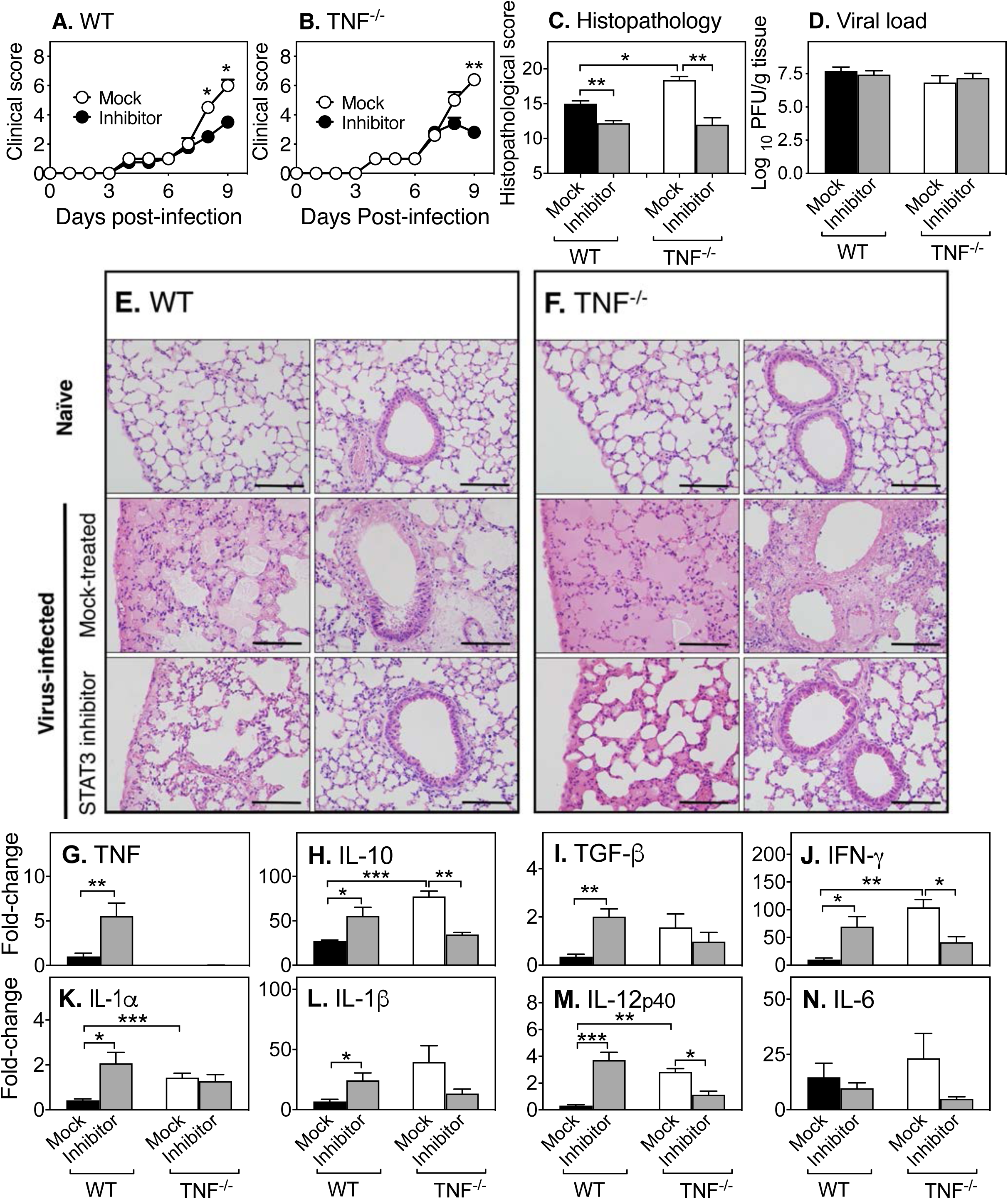
STAT3 inhibition dampens ECTV infection induced lung pathology and modulates inflammatory cytokine responses but does not affect virus load. Groups of 5 WT and 5 TNF^-/-^ ECTV-infected mice were treated on days 7 and 8 p.i. with 5 mg/kg of the STAT3 inhibitor, S3I-201, and mock-treated groups were given inhibitor diluent. Clinical scores of (A) WT and (B) TNF^-/-^ mice based on the clinical scoring criteria (Table S1). (C) Lung histopathological scores and (D) viral load at day 9 p.i. (E, F). Representative lung sections are shown at 400x magnification and bars correspond to 100 µm. Levels of expression of mRNA transcripts for (G) TNF, (H) IL-10, (I) TGF-β, (J) IFN-γ, (K) IL-1α, (L) IL-1β, (M) IL-12p40 and (N) IL-6. Data are expressed as means ± SEM and analyzed using Mann-Whitney tests for (A)-(C). For (D) data were log transformed and analyzed by two-way ANOVA, with Holm-Sidak’s correction for multiple comparison and the broken line corresponds to the limit of virus detection. For (G)-(N), data were analyzed by two-way ANOVA with Holm-Sidak’s correction for multiple comparison. Results are representative of 2 separate experiments.

Treatment with S3I-201 affected the levels of expression of a number of inflammatory cytokines. In WT animals, which are able to recover from infection without significant lung pathology, inhibitor treatment resulted in significant increases in the levels of mRNA for TNF (Figure 6G), IL-10 (Figure 6H), TGF-β (Figure 6I), IFN-γ (Figure 6J), IL-1α (Figure 6K), IL-1β (Figure 6L), and IL-12p40 (Figure 6M). In sharp contrast, in TNF^-/-^ mice, which exhibit severe lung pathology and succumb to infection, treatment with the STAT3 inhibitor resulted in a significant reduction in the levels of some cytokine mRNA transcripts compared to the mock-treated group. A reduction in mRNA levels was observed for IL-10 (Figure 6H), IFN-γ (Figure 6J), and IL-12p40 (Figure 6M). Although levels of mRNA for IL-1β (Figure 6L) and IL-6 (Figure 6N) were also reduced compared to the mock-treated group, they were not statistically significant. Crucially, STAT3 inhibitor treatment reduced mRNA levels of a number of cytokines in TNF^-/-^ mice down to those observed in mock-treated WT animals. Therefore, the effect of STAT3 inhibition on the expression of mRNA transcripts for inflammatory cytokines is dictated by the presence or absence of TNF.

### TNF^-/-^ mice recover from ECTV infection after extended treatment with anti-IL-6 or TGF-β

Short-term treatment with anti-cytokine mAb or STAT3 inhibitor significantly reduced lung pathology in TNF^-/-^ animals. It was thus of interest to determine whether prolonged treatment with these agents would allow TNF^-/-^ mice to recover from an otherwise lethal infection. ECTV-infected TNF^-/-^ mice were treated with either a control mAb or mAb against IL-6, IL-10R or TGF-β beginning at day 7 p.i., and every 2 days thereafter until day 20 p.i. A separate ECTV-infected group was treated with the STAT3 inhibitor from day 7 p.i. and every day thereafter until day 20 p.i. All animals were monitored for 22 days. For virus load and histopathological scores, organs were collected from animals that were moribund (and euthanized for ethical reasons) or found dead before day 22 p.i., and those that were sacrificed on day 22. Therefore, statistical analysis was not performed for virus load and histopathological scores.

Control mAb- and anti-IL-10R-treated mice exhibited 100% mortality by days 10-13 p.i., whereas animals treated with anti-IL-6 or anti-TGF-β survived the infection (Figure 7A). Four of 5 animals treated with the STAT3 inhibitor died between days 11-18, and one animal survived until the day of sacrifice. Extended treatment with anti-IL-10R did not improve weight loss whereas all animals treated with anti-IL-6 or anti-TGF-β showed improvements in body weights from day 13 (Figure 7B) and clinical scores (Figure 7C) within 1-2 days after initiation of treatment. The one STAT3 inhibitor-treated mouse that survived gained weight from day 19 p.i. (Figure 7B) in parallel with a reduction in clinical scores (Figure 7C). Viral load in lungs of control mAb-, anti-IL-10R- or STAT3 inhibitor-treated mice were very high compared to titers in mice treated with anti-IL-6 or anti-TGF-β, which were close to the limit of detection (Figure 7D). Viral load in the one mouse treated with the STAT3 inhibitor that survived was also below the limit of detection.

**Figure 7.**
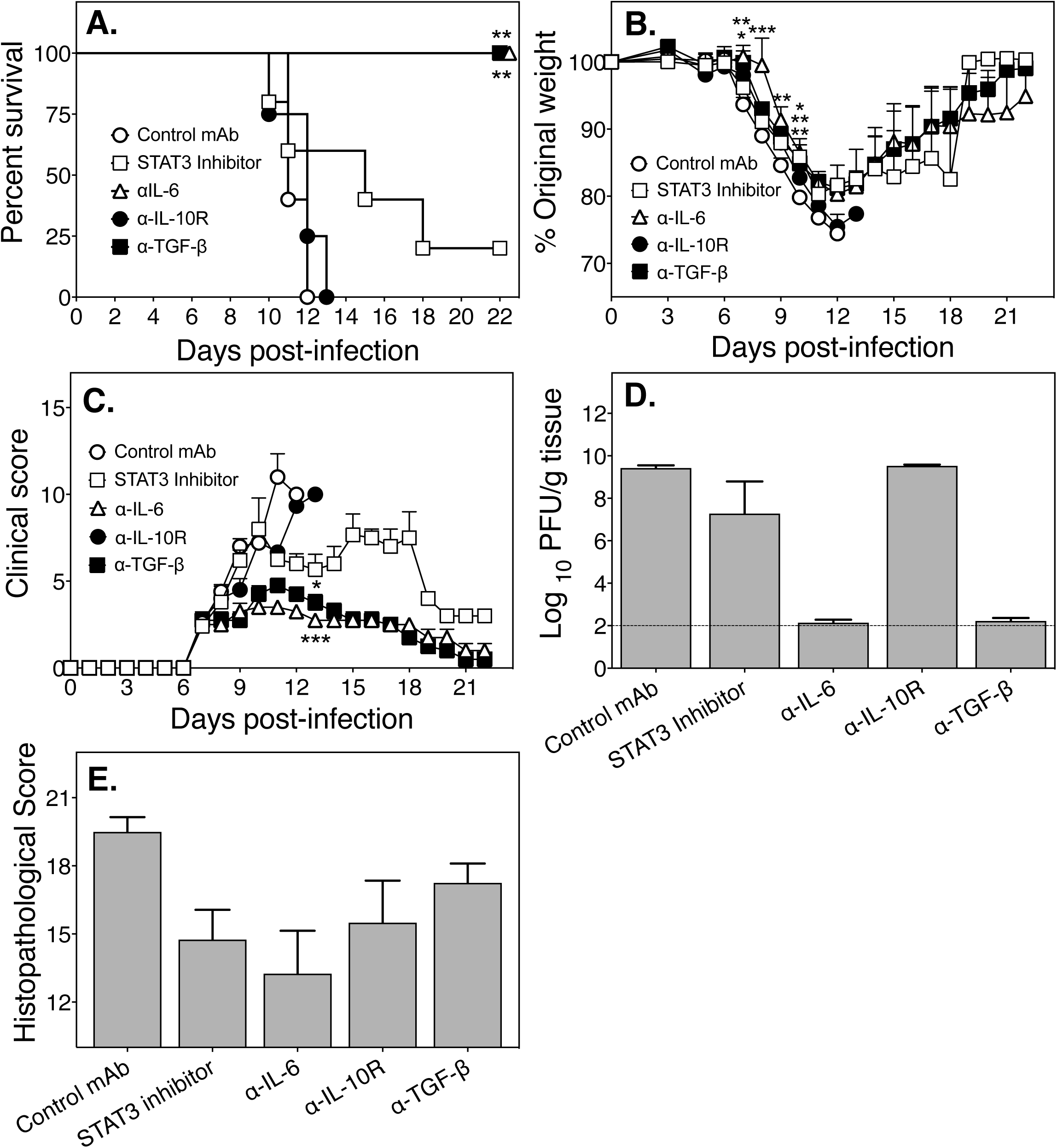
Long-term treatment with anti-IL-6 or anti-TGF-β protects TNF^-/-^ mice from lethal disease. A group of 5 TNF^-/-^ ECTV-infected mice were treated every day from days 7-21 p.i. with 5 mg/kg STAT3 inhibitor and other groups were treated with control rat IgG mAb, anti-IL-6, anti-IL-10R, or anti-TGF-β at 500 µg i.p. every 2 days from day 7 until day 21 p.i., and monitored until day 22. (A) Survival, (B) weights and (C) clinical scores were monitored during the course of infection. (D) Lung viral load and (E) histopathological scores of lung H&E sections for organs collected from animals that were moribund and euthanized for ethical reasons or found dead before day 22 p.i., and those that were sacrificed on day 22. Data are expressed as means ± SEM. For (A), (B) and (C), data for STAT3 inhibitor- or anti-cytokine mAb-treated mice were compared with control mAb-treated mice for each day. Survival curves (A) were compared using Log-rank (Mantel-Cox) statistical test. Weights (B) were compared by multiple unpaired t-tests and the Holm-Sidak’s correction for multiple comparisons. Clinical scores (C) were compared by the two-tailed Mann-Whitney test. For (D), viral load data were log transformed and the broken line corresponds to the limit of detection in the plaque assay. Statistical analysis was not performed for viral load and histopathological scores. Results are representative of 2 separate experiments.

Histologically, only prolonged treatment with anti-IL-6 reduced lung pathology (Figure 7E). All control mAb-, anti-TGF-β- or anti-IL-10R-treated mice and 4 of 5 STAT3 inhibitor-treated mice had significant lung pathology as revealed by the histopathological scores (Figure 7E). The finding with anti-TGF-β treatment was surprising as even with this level of pathology, the viral load was close to the limit of detection, the clinical scores were close to baseline and the animals regained their loss in body weights.

The results indicate that the immune dysregulation and pulmonary pathology in TNF^-/-^ mice following infection with ECTV were due to excessive production of some inflammatory and regulatory cytokines and that blockade of those cytokines can protect mice from an otherwise lethal infection.

## DISCUSSION

TNF plays fundamental roles in homeostasis, acute inflammation and antimicrobial defense but excessive or chronic production of the cytokine can cause serious pathology and mortality. Paradoxically, our results have shown that TNF deficiency in mice also results in significant pathology and death during mousepox, a surrogate model for smallpox. We also established that mTNF alone was necessary and sufficient to protect against lung pathology in ECTV infected mice. The immunopathology was not due to excessive leukocyte infiltration into the lungs as numbers and phenotypes of leukocytes present in virus-infected WT and TNF^-/-^ groups were not different. The numbers and magnitude of the cytolytic activities of two key cell populations, i.e., NK and CD8^+^ T cells, present in the lungs and spleens of infected animals were not also affected in the absence of TNF. Both NK cell and CD8^+^ T cell-mediated CTL responses are essential for early virus control and recovery of mice from ECTV infection (Chaudhri et al., 2006; Chaudhri et al., 2004; Delano and Brownstein, 1995; Fang et al., 2008; Fang and Sigal, 2005; Karupiah et al., 1996; Parker et al., 2007) and TNF can influence proliferation and effector functions of CD8^+^ T cells (Damjanovic et al., 2011; Peper and Van Campen, 1995; Scheurich et al., 1987). The absence of any impact of TNF deficiency on NK cell and CTL responses is also consistent with a lack of any effect on viral load since deficiencies in either effector cell type can significantly augment viral load.

The antimicrobial activity of TNF is well-documented for bacterial, fungal and some viral pathogens. In humans, treatment with TNF blocking agents can exacerbate some viral infections (Murdaca et al., 2015; Vigano et al., 2012) but not others like influenza A (Shale et al., 2010). In mice, TNF deficiency did not have any effects on influenza A virus titers (Damjanovic et al., 2011; Peper and Van Campen, 1995), but exacerbated illness and heightened lung immunopathology. Our results with the ECTV model are consistent with findings in the influenza A virus model. The current study indicates that TNF, unlike IFN-α/β and IFN-γ (Chaudhri et al., 2004; Karupiah et al., 1993; Panchanathan et al., 2005), has no antiviral activity against ECTV in C57BL/6 mice but is necessary for regulation of inflammation.

WT C57BL/6 mice produced high levels of TNF during the resolution stage (day 12 p.i.) of ECTV infection. In the absence of TNF, levels of IFN-γ, IL-6, IL-10 and TGF-β were significantly higher than in WT mice. Both IL-6 and IFN-γ are considered pro-inflammatory cytokines with the capacity to cause tissue damage (Rincon, 2012; Schroder et al., 2004). On the other hand, TGF-β and IL-10 are known to regulate the immune response but they can also mediate pro-inflammatory activities (Sanjabi et al., 2009). IL-10 can regulate immunity by limiting inflammation to prevent tissue damage but high levels of the cytokine can itself cause chronic inflammation and immunopathology (Couper et al., 2008). The short-term cytokine neutralization experiments provided direct evidence that the exaggerated lung tissue destruction in the absence of TNF was due to dysregulated production of IFN-γ, IL-6, IL-10 and TGF-β. The timing of cytokine neutralization *in vivo, i.e.* co-incident with generation of pulmonary edema and lung pathology, was crucial as some of these cytokines also play very important protective roles during the early stages of the infection. In particular, IFN-γ and IL-6 are critical for ECTV control very early during the course of infection (Chaudhri et al., 2004; Ivashkiv, 2010; Karupiah et al., 1993; O’Gorman et al., 2010). TNF thus plays a critical role in the suppression of immunopathology at the resolution stage of infection through regulation of inflammatory/ regulatory cytokine production.

The finding that dampening of IFN-γ, IL-6, IL-10 or TGF-β reduced lung pathology suggested at least two, non-mutually exclusive, possibilities. First, IFN-γ, IL-6, IL-10 and TGF-β can phosphorylate and activate STAT3 (McGeachy et al., 2007; Niemand et al., 2003; Qing and Stark, 2004; Yamamoto et al., 2001; Zhong et al., 1994), potentially involving a common cytokine-signaling pathway. Second, the different signaling pathways activated by each of these cytokines can cross-regulate each other. For example, STAT3 and NF-κB can regulate the expression of inflammatory genes in a cooperative manner (Oeckinghaus et al., 2011). TNF can activate STAT3 by induction of IL-6 through NF-κB activation (Zhong et al., 1994). However, TNF can also inhibit IL-6-mediated STAT3 activation in macrophages through the recruitment of SOCS3 (Bode et al., 2003). In our present study, the anti-inflammatory role of TNF during ECTV infection was evidenced by low levels of PIAS3 in the lungs of TNF^-/-^ mice compared to the WT animals.

PIAS3 not only negatively regulates STAT3 but may also physically interact with p65 subunit of NF-κB, thereby inhibiting the latter’s activity (Jang et al., 2004). PIAS3 is a small ubiquitin-like modifier (SUMO) E3 ligase that belongs to a family of STAT signaling regulators, and the main cellular inhibitor of STAT3 (Chung et al., 1997). Levels of PIAS3 expression are directly correlated with inhibition of STAT3 DNA-binding and transcriptional activity (Dabir et al., 2009). Hence, the increased levels of pSTAT3 in the anti-cytokine mAb-treated mice was likely inhibited by the correspondingly increased levels of PIAS3. IL-6 blockade also induced high levels of SOCS3 and the combined actions of PIAS3 and SOCS3 in this group of animals likely contributed to effective control of lung pathology and viral load.

Dysregulated or prolonged activation of STAT3 can lead to severe pathologic outcomes and disease (Grivennikov and Karin, 2008). Our data indicates that overactivation of the STAT3 signaling pathway, at least in part, contributed to the exacerbated immunopathology in TNF-deficient mice and that lung pathology could be ameliorated by short-term inhibition of STAT3 activation. Nonetheless, long-term treatment with the inhibitor was ineffective in reducing lung pathology or overcoming morbidity and mortality. Such an outcome is likely due to the possibility that some cytokines or factors produced through the STAT3 signaling pathway may be directly or indirectly essential for down-regulating inflammation or involved in lung tissue repair during the resolution phase of the infection. A number of cytokine transcripts, including TGF-β, were down-regulated by STAT3 inhibitor treatment in TNF^-/-^ mice. TGF-β is critical for tissue repair and although long-term treatment of TNF^-/-^ mice with anti-TGF-β protected ECTV-infected mice from death, the lung pathology was not resolved fully at day 22 p.i. Additionally, prolonged anti-IL-10 treatment, unlike short-term treatment, was not protective as this cytokine is also important for regulation of the inflammatory response.

The outbred human population exhibited varying degrees of susceptibility to smallpox (Fenner, 1988) and similarly, inbred strains of mice are either resistant or susceptible to mousepox. ECTV-resistant mice like the C57BL/6 strain generate a strong inflammatory response and produce high levels of IL-2, IFN-γ and TNF very early in infection (Chaudhri et al., 2004). In these animals, a polarized type I cytokine response is not only closely associated with induction of robust cell-mediated immunity but also generation of potent antibody responses, both of which are critical for recovery from infection (Chaudhri et al., 2006; Fang and Sigal, 2005). In contrast, ECTV-susceptible strains such as BALB/c mice produce significantly lower levels of these cytokines, associated with very weak inflammatory responses and cell-mediated immunity (Chaudhri et al., 2004). In the current study, we have found that TNF deficiency in ECTV-resistant mice results in significant lung pathology and death during a respiratory ECTV infection. The increased susceptibility of mice was clearly not due to an increase in viral load. These results might appear to contradict with a recent report that TNF plays an important antiviral role in the recovery of the ECTV-susceptible BALB/c mice (Alejo et al., 2018). In that study, infection of the BALB/c strain with the CrmD deletion mutant virus augmented inflammation and cell-mediated immunity, resulting in effective virus control and complete recovery from an otherwise lethal infection. There are at least two reasons for the observed differences in the outcomes of infection with the the mutant virus in the two strains of mice and these are discussed below.

First, the potent immune response generated by C57BL/6 mice can largely overcome the effects of the various host response modifiers that ECTV encodes, including CrmD. C57BL/6 mice produce significantly higher levels of TNF than BALB/c mice in response to WT ECTV infection (Chaudhri et al., 2004). Our prediction is that in C57BL/6 mice infected with the CrmD deletion mutant virus, we would expect to see further increases in TNF levels. That is indeed the case as C57BL/6 mice develop significant lung pathology due to excessive TNF production, an overexuberant inflammatory response in the lung and succumb to mousepox (see accompanying manuscript). Because BALB/c mice produce low levels of TNF, CrmD is likely to have neutralized the cytokine, resulting in poor quality of the inflammatory and cell-mediated immune responses (Alejo et al., 2018). Second, we used the i.n. route of inoculation in our studies with C57BL/6 mice whereas Alcami’s group (Alejo et al., 2018) used the subcutaneous route. ECTV can be naturally transmitted via the subcutaneous route through scratches or bites from infected animals as well as via the respiratory route through aerosols. Both routes of virus transmission result in systemic infection, but the i.n. route is more sensitive, with virus replicating to high titers in the lungs, and makes the resistant C57BL/6 strain succumb to mousepox at low doses of virus. Curiously, we did not see any significant differences in the susceptibility of WT and TNF^-/-^ C57BL/6 mice inoculated with varying doses of ECTV through the subcutaneous route (data not shown). That finding suggested that a requirement for TNF to protect against ECTV infection in C57BL/6 mice, specifically to regulate the local inflammatory response, depends on the route through which virus is transmitted to the host.

In summary, TNF has no direct antiviral activity against ECTV but is necessary to regulate lung inflammation. Respiratory ECTV infection of TNF^-/-^ mice caused lethal lung pathology due to dysregulation and excessive production of some specific cytokines and overactivation of STAT3. Short-term blockade of any of these cytokines with mAb or inhibition of STAT3 activation ameliorated lung pathology in ECTV-infected mice. Our data indicates that targeting specific cytokines or cytokine signaling pathways to reduce or ameliorate lung inflammation during a viral infection is possible but that the timing and duration of the interventive measure will be critical.

## Acknowledgements

This work was supported by a grant from the National Health and Medical Research Council of Australia to G.K. and G.C. (grant ID APP 1007980). The funders had no role in study design, data collection and interpretation, or the decision to submit the work for publication. We would like to thank Professor Jane Dahlstrom of the Canberra Hospital, Australian Capital Territory, Australia for helping us to develop the scoring system for the histopathology of lung sections. We also thank Dr. Jonathon D. Sedgwick, Boehringer Ingelheim Pharmaceuticals Inc., Ridgefield, USA. We gratefully acknowledge Professor Antonio Alcami, Centro de Biología Molecular Severo Ochoa, Madrid for the gift of the Naval strain of WT and CrmD mutant ECTV and the assistance of staff at the Australian National University Phenomics Facility for animal breeding and the John Curtin School of Medical Research Microscopy and Flow Cytometry Research Facility.

## Author contributions

Conceptualization, M.J.T.K., E.N, G.C. and G.K.; Methodology, M.J.T.K., E.N., Z.A.R., P.P., G.C. and G.K.; Investigation, M.J.T.K., E.N., Z.A.R., T.P.N.; Writing – Original Draft, M.J.T.K., G.C. and G.K.; Writing – Review & Editing, M.J.T.K., E.N, H.K., T.P.N., G.C. and G.K.; Funding Acquisition, G.C. and G.K.; Resources, S.R.R., H.K., G.C. and G.K.; Supervision, G.C. and G.K.

## Methods

### Mice

Six- to twelve-week old, female WT, TNF^-/-^ (Korner et al., 1997), mTNF^Δ/Δ^ (Ruuls et al., 2001), IFN-α/βR^-/-^ (Muller et al., 1994) and IFN-γ^-/-^ (Dalton et al., 1993) mice on a C57BL/6 background were bred under specific pathogen-free conditions at the Australian Phenomics Facility, Australian National University, Canberra, Australia. Animal experiments were performed in accordance with protocols approved by the ANU Animal Ethics and Experimentation Committee (Protocol numbers A2011/011 and A2014/018).

### Cell lines and viruses

BS-C-1 cells (ATCC No. CCL-26), MC57G (ATCC No. CRL-2295) and YAC-1 (ATCC No. TIB-160) were cultured in Eagle’s minimum essential medium (EMEM) supplemented with 2 mM L-glutamine (Sigma Aldrich), antibiotics (penicillin, 120 μg/mL, streptomycin, 200 μg/mL and neomycin sulphate), 1 mM 4-(2-hydroxyethyl)-1-piperazineethanesulfonic acid (HEPES) and 10% fetal calf serum (FCS).

The Moscow strain of ECTV (ATCC No. VR-1374) was used to infect mice in all experiments except for data shown in Figure 1H, for which the Naval strain of WT ECTV (ECTV^WT^) and the CrmD deletion mutant (ECTV^ΔCrmD^) (Alejo et al., 2018) were used. Viruses were propagated in BS-C-1 cells, semi-purified using a sucrose cushion and single-use aliquots were stored at −80°C as described elsewhere in detail (Chaudhri et al., 2018).

### Virus infection, animal weights, clinical scores and treatments

Tribromoethanol-anesthetized mice (160-240 mg/kg i.p.) were infected through the i.n. route with 25 PFU of ECTV in 30 μL PBS. Clinical manifestation of disease during ECTV infection was assessed using the scoring system (Table S1) with 5 clinical parameters. In some experiments, mice were administered i.p. with STAT3 inhibitor VI, S3I-201 (Merck Millipore cat. no. 573102) at 5 mg/kg, or neutralizing/blocking mAb against IFN-γ (R4.6A2, rat IgG_1_, cat. no. BE0054), IL-6 (clone MP5-20F3, rat IgG_1,_ cat. no. BE0046), IL-10R (1B1.3A, rat IgG_1_, cat. no. BE0050), TGF-β (1D11.16.18, mouse IgG_1_, cat. no. BE0057) or isotype control mAb (HRPN, rat IgG_1_, cat. no. BE0088) at 500 μg on day 7 p.i. Purified mAb were purchased from Bio X Cell. Mice were individually ear-tagged, weighed and clinical scores recorded daily on a scale from 0 to 3 for each criterion, with a maximum score of 15. For ethical reasons, mice that were severely moribund and/or lost ≥ 25% of the original body weight were euthanized and considered dead the following day.

### Histology and microscopic assessment of lung pathology

The left lobe of lung tissue was removed and fixed in 10% neutral-buffered formaldehyde at room temperature for 24 h, embedded in paraffin, cut in 6 μm sagittal sections and stained with hematoxylin and eosin (H&E) for analysis with a bright field microscope (Olympus IX 71). A semi-quantitative scoring system for lung pathology was developed (Table S2) with 6 pathologic parameters. Individual slides were scored in a single-blinded fashion on a scale from 0 to 4 for each criterion, with a maximum score of 24.

### Lung total protein concentration and wet-to-dry weight ratio

Total protein concentration in bronchoalveolar lavage fluid (BALF) were measured by the Bradford Assay using Protein Assay Dye Reagent Concentrate (Bio-Rad) according to the manufacturer’s protocol. For the quantification of pulmonary edema, the left lung lobe was removed and immediately wrapped in a pre-weighed aluminium foil, weighed (wet lung weight) and dried in an oven at 80°C for 72 h. The weights (oven-dried lung weight) of samples were recorded after the 72-h period to determine wet-to-dry weight ratio (wet lung weight / oven-dried lung weight).

### Flow cytometric analysis of lung leukocytes

Individual lung tissue samples were cut into small pieces and digested in EMEM containing 4 μg DNAse I, grade II and 4 mg collagenase A (Roche Applied Science) and placed in a 37°C water bath for 1 h. Undigested tissue was further processed by passing through a 19-G needle attached to a syringe and then filtered through a 70 μm nylon mesh. Cells were washed with Hank’s balanced salt solution and red blood cells lysed with red cell lysis buffer (150 mM NH_4_Cl, 10 mM KHCO_3_, 0.1 mM Na_2_EDTA, pH 7.4) for 1 min. Cells were next washed and resuspended in PBS containing 2% FCS. For labelling, FcγII/III receptors (FcγR) were first blocked with anti-FcγR mAb (clone 2.4G2), the cells were washed, and stained with CD45.2-FITC (to identify leukocytes) and cell subset specific fluorochrome-conjugated mAb (all were obtained from BD Biosciences and Biolegend). Data were acquired on a BD LSR Fortessa flow cytometer with BD FACS Diva software and analyzed using FlowJo Software version 9.5 (Tree Star, Inc).

### Determination of cytokine protein levels

The levels of IFN-γ, IL-6, IL-10 and TNF in the lung homogenates were measured using the Mouse Inflammation Cytokine Bead Array Kit (BD Biosciences) according the manufacturer’s protocol. Acquisition of data was done using a BD FACSCalibur flow cytometer with BD Cell Quest software. Data were analyzed using FlowJo Software version 9.5. TGF-β levels were measured using capture ELISA kits (Biolegend), performed according to the manufacturer’s protocol. Optical density was measured at 450 nm with SOFTmax Pro software (Molecular Devices Corp).

### SDS PAGE and Western blot analysis

Lung homogenates and a pre-stained recombinant protein ladder (Bio-Rad) were run in a 12% SDS-polyacrylamide gel and were transferred to a nitrocellulose membrane (Bio-Rad). After blocking with 1 M glycine, 5% w/v dry skim milk blocking solution, the membrane was incubated with primary antibody (STAT3, polyclonal rabbit IgG, cat. no. 9132; pSTAT3 (Tyr705), polyclonal rabbit IgG, cat. no. 9131; PIAS3, polyclonal rabbit IgG, cat. no. 4164, all from Cell Signaling Technology; or SOCS3, Abcam, cat. no. ab16030). The immunoreactive proteins were visualized with horseradish peroxidase (HRP)-conjugated secondary antibody (polyclonal goat anti-rabbit IgG, Santa Cruz Biotechnology, cat.no. sc-2004) and SuperSignal West Pico Chemiluminescent Substrate (Thermo Fisher Scientific) in a chemiluminescent imager (ImageQuant LAS 4000, GE Healthcare Life Sciences).

### Immunohistochemistry of lung sections

Immunohistochemistry was performed on formaldehyde-fixed and paraffin-embedded lung sections. Samples were deparaffinized and rehydrated through different concentrations of xylene and ethanol, followed by antigen retrieval in Tris/EDTA Buffer. The sections were incubated in primary TNF antibody (MP6-XT3, rat IgG_1_) and secondary HRP-conjugated goat anti-rat IgG (Santa Cruz Biotechnology, cat. no. sc-52619 and sc-2006, respectively). The reaction was developed using a 3,3’-diaminobenzidine substrate buffer and counterstained with H&E for analysis with a bright field microscope (Olympus IX 71).

### RNA extraction

Lung tissue from individual mice were homogenized in 1 mL of TRIsure reagent with TissueLyser II (Qiagen, cat. no. 853000) and as per the manufacturer’s instructions and samples were centrifuged for 15 min at 12,000 x g at 4°C to remove cellular debris. In the final step, RNA pellets were air-dried and resuspended in 30-50 µL RNAse-free water and incubated for 10 min at 60°C to ensure they were dissolved. RNA concentration was determined using a Nanodrop ND-1000 spectrophotometer. Samples were stored at −80°C until cDNA synthesis.

### cDNA generation

Tetro cDNA Synthesis Kit (Bioline, cat. no. BIO-65043) was used to synthesize single-stranded cDNA from 1 µg RNA in 12 µL RNAse-free water following the manufacturer’s instructions. Briefly, the priming premix was prepared on ice by mixing Oligo (dT)_18_ primer mix to a final concentration of 0.5 µg/µL, 10mM dNTP mix, 5x RT Buffer, 10 units/µL RiboSafe RNase Inhibitor and Tetro Reverse Transcriptase (200 unit/μL). To each sample, 8 µl of the mastermix was added and incubated at 45°C for 30 min. The reaction was terminated by incubating samples at 85°C for 5 min then chilled on ice. Samples were diluted with 180 µL RNAse free water and stored at −20°C until further use.

### Quantitative reverse transcription real-time polymerase chain reaction (qRT-PCR)

For each gene of interest, a real-time PCR reaction mixture was prepared containing 10 µL of SsoAdvanced™ SYBR® Green Supermix (Bio-Rad, cat. no. 172-5265), 1 µL each of 5 µM gene-specific forward and reverse primer pair, and 6 µL of RNAse-free water. 2 µL of the appropriate cDNA template and 18 µL of the reaction mixture were added to the wells of iQ 96-well semi-skirted PCR Plate (Bio-Rad, cat. no. 223-9441). Amplification was achieved using Bio-Rad iQ5 Real-time PCR Detection System. Relative changes in gene expression was determined using delta/delta cycle threshold (2^-ΔΔCT^) analysis method (Livak and Schmittgen, 2001). *Ubiquitin C* (*UBC*) was used as the internal control. Results are reported as the fold difference relative to a calibrator cDNA (naïve mice) prepared in parallel with the experimental cDNAs.

### ^51^Chromium (^51^Cr) release cytotoxic assays

CTL and NK cell responses were measured by the ^51^Cr release assay as described previously (Karupiah et al., 1990; Karupiah et al., 1991). Briefly, for CTL assays, 6 x 10^6^ MC57G target cells that were either uninfected or infected with ECTV (10 PFU/cell) were labelled with 20 μCi ^51^Cr (Na_2_^51^CrO_4_) and incubated in a humidified atmosphere of 5% CO_2_ and 95% air for 1 h. Cells were washed three times and resuspended in complete medium and 2 x 10^4^ cells were added into wells of U-bottom 96-well plates. Effector cells (single cell suspensions of digested lungs or splenocytes) were added to wells with target cells in triplicate to obtain effector cell:target cell ratios of 100:1, 33:1, 11:1 and 3.7:1. Control wells included target cells only (spontaneous release) or target cells plus 1% Triton X (maximum release). After incubation for 6 h in a humidified atmosphere of 5% CO_2_ and 95% air, the 96-well plates were centrifuged at 300 x *g* for 5 min, and 25 μl of supernatant from each well was transferred to a corresponding well in Luma-96 plate (Perkin-Elmer, Boston, USA). Plates were air dried overnight and radioactive counts per minute (cpm) were measured using a TopCount NXT^TM^ microplate scintillation counter (Packard Bioscience Company, Meriden CT, USA). Percent specific lysis was calculated from the mean cpm of triplicate wells, using the following equation:

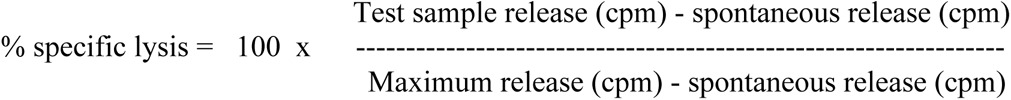

Virus-specific CTL activity was determined by subtracting % specific lysis values of uninfected targets from infected targets.

For NK cell responses, YAC-1 target cells were used without the need for infection with ECTV. YAC-1 cells were labelled with ^51^Cr as described for MC57G cells. Effector and target cells were incubated for 4 h after which supernatants were collected to determine cpm as described for the CTL assay.

### Statistical analysis

Statistical analyses of experimental data, as indicated, were performed using GraphPad Prism 8 (GraphPad Software, Inc.). A p value of <0.05 was taken to be significant, *, p ≤ 0.05; **, p ≤ 0.01; ***, p ≤ 0.001; ****, p ≤ 0.0001. Log-rank (Mantel-Cox) test, two-tailed Mann-Whitney test, unpaired t-test with Holm-Sidak’s correction and two-way ANOVA with Holm-Sidak’s correction for multiple comparison were used to compare results.

## REFERENCES

1. Alcami, A. (2003). Viral mimicry of cytokines, chemokines and their receptors. Nat Rev Immunol. 3(1), 36–50.

2. Alejo, A., Ruiz-Arguello, M.B., Pontejo, S.M., Fernandez de Marco, M.D.M., Saraiva, M., Hernaez, B., and Alcami, A. (2018). Chemokines cooperate with TNF to provide protective anti-viral immunity and to enhance inflammation. Nat Commun. 9(1), 1790.

3. Alejo, A., Saraiva, M., Ruiz-Arguello, M.B., Viejo-Borbolla, A., de Marco, M.F., Salguero, F.J., and Alcami, A. (2009). A method for the generation of ectromelia virus (ECTV) recombinants: in vivo analysis of ECTV vCD30 deletion mutants. PLoS One. 4(4), e5175.

4. Arnett, H.A., Mason, J., Marino, M., Suzuki, K., Matsushima, G.K., and Ting, J.P. (2001). TNF alpha promotes proliferation of oligodendrocyte progenitors and remyelination. Nat Neurosci. 4(11), 1116–1122.

5. Babon, J.J., Varghese, L.N., and Nicola, N.A. (2014). Inhibition of IL-6 family cytokines by SOCS3. Semin Immunol. 26(1), 13–19.

6. Bean, A.G., Roach, D.R., Briscoe, H., France, M.P., Korner, H., Sedgwick, J.D., and Britton, W.J. (1999). Structural deficiencies in granuloma formation in TNF gene-targeted mice underlie the heightened susceptibility to aerosol Mycobacterium tuberculosis infection, which is not compensated for by lymphotoxin. J Immunol. 162(6), 3504–3511.

7. Belisle, S.E., Tisoncik, J.R., Korth, M.J., Carter, V.S., Proll, S.C., Swayne, D.E., Pantin-Jackwood, M., Tumpey, T.M., and Katze, M.G. (2010). Genomic profiling of tumor necrosis factor alpha (TNF-alpha) receptor and interleukin-1 receptor knockout mice reveals a link between TNF-alpha signaling and increased severity of 1918 pandemic influenza virus infection. J Virol. 84(24), 12576–12588.

8. Black, R.A., Rauch, C.T., Kozlosky, C.J., Peschon, J.J., Slack, J.L., Wolfson, M.F., Castner, B.J., Stocking, K.L., Reddy, P., Srinivasan, S., et al. (1997). A metalloproteinase disintegrin that releases tumour-necrosis factor-α from cells. Nature. 385, 729.

9. Bode, J.G., Schweigart, J., Kehrmann, J., Ehlting, C., Schaper, F., Heinrich, P.C., and Haussinger, D. (2003). TNF-alpha induces tyrosine phosphorylation and recruitment of the Src homology protein-tyrosine phosphatase 2 to the gp130 signal-transducing subunit of the IL-6 receptor complex. J Immunol. 171(1), 257–266.

10. Carow, B., and Rottenberg, M.E. (2014). SOCS3, a Major Regulator of Infection and Inflammation. Front Immunol. 5, 58.

11. Chaudhri, G., Kaladimou, G., Pandey, P., and Karupiah, G. (2018). Propagation and Purification of Ectromelia Virus. Curr Protoc Microbiol. 51(1), e65.

12. Chaudhri, G., Panchanathan, V., Bluethmann, H., and Karupiah, G. (2006). Obligatory requirement for antibody in recovery from a primary poxvirus infection. J Virol. 80(13), 6339–6344.

13. Chaudhri, G., Panchanathan, V., Buller, R.M., van den Eertwegh, A.J., Claassen, E., Zhou, J., de Chazal, R., Laman, J.D., and Karupiah, G. (2004). Polarized type 1 cytokine response and cell-mediated immunity determine genetic resistance to mousepox. Proc Natl Acad Sci U S A. 101(24), 9057–9062.

14. Chung, C.D., Liao, J., Liu, B., Rao, X., Jay, P., Berta, P., and Shuai, K. (1997). Specific Inhibition of Stat3 Signal Transduction by PIAS3. Science. 278(5344), 1803–1805.

15. Couper, K.N., Blount, D.G., and Riley, E.M. (2008). IL-10: the master regulator of immunity to infection. J Immunol. 180(9), 5771–5777.

16. Dabir, S., Kluge, A., and Dowlati, A. (2009). The Association and Nuclear Translocation of the PIAS3-STAT3 Complex Is Ligand and Time Dependent. Mol Cancer Res. 7(11), 1854–1860.

17. Dalton, D.K., Pitts-Meek, S., Keshav, S., Figari, I.S., Bradley, A., and Stewart, T.A. (1993). Multiple defects of immune cell function in mice with disrupted interferon-gamma genes. Science. 259(5102), 1739–1742.

18. Damjanovic, D., Divangahi, M., Kugathasan, K., Small, C.L., Zganiacz, A., Brown, E.G., Hogaboam, C.M., Gauldie, J., and Xing, Z. (2011). Negative regulation of lung inflammation and immunopathology by TNF-alpha during acute influenza infection. Am J Pathol. 179(6), 2963–2976.

19. Delano, M.L., and Brownstein, D.G. (1995). Innate resistance to lethal mousepox is genetically linked to the NK gene complex on chromosome 6 and correlates with early restriction of virus replication by cells with an NK phenotype. J Virol. 69(9), 5875–5877.

20. Domm, S., Cinatl, J., and Mrowietz, U. (2008). The impact of treatment with tumour necrosis factor-alpha antagonists on the course of chronic viral infections: a review of the literature. Br J Dermatol. 159(6), 1217–1228.

21. Fang, M., Lanier, L.L., and Sigal, L.J. (2008). A role for NKG2D in NK cell-mediated resistance to poxvirus disease. PLoS Pathog. 4(2), e30.

22. Fang, M., and Sigal, L.J. (2005). Antibodies and CD8+ T cells are complementary and essential for natural resistance to a highly lethal cytopathic virus. J Immunol. 175(10), 6829–6836.

23. Fenner, F. (1988). Smallpox and its eradication (World Health Organization: Geneva).

24. Garcia-Gonzalez, E., Guidelli, G.M., Bardelli, M., and Maggio, R. (2012). Mucocutaneous leishmaniasis in a patient treated with anti-TNF-α therapy. Rheumatology. 51(8), 1517–1518.

25. Goldstein, I., Paakinaho, V., Baek, S., Sung, M.H., and Hager, G.L. (2017). Synergistic gene expression during the acute phase response is characterized by transcription factor assisted loading. Nat Commun. 8(1), 1849.

26. Grivennikov, S., and Karin, M. (2008). Autocrine IL-6 signaling: a key event in tumorigenesis? Cancer Cell. 13(1), 7–9.

27. Herbein, G., and O’Brien, W.A. (2000). Tumor necrosis factor (TNF)-alpha and TNF receptors in viral pathogenesis. Proc Soc Exp Biol Med. 223(3), 241–257.

28. Hillmer, E.J., Zhang, H., Li, H.S., and Watowich, S.S. (2016). STAT3 signaling in immunity. Cytokine Growth Factor Rev. 31, 1–15.

29. Ivashkiv, L.B. (2010). STAT activation during viral infection in vivo: where’s the interferon? Cell Host Microbe. 8(2), 132–135.

30. Jang, H.D., Yoon, K., Shin, Y.J., Kim, J., and Lee, S.Y. (2004). PIAS3 suppresses NF-kappaB-mediated transcription by interacting with the p65/RelA subunit. J Biol Chem. 279(23), 24873–24880.

31. Kalliolias, G.D., and Ivashkiv, L.B. (2016). TNF biology, pathogenic mechanisms and emerging therapeutic strategies. Nat Rev Rheumatol. 12(1), 49–62.

32. Karupiah, G., Buller, R.M., Van Rooijen, N., Duarte, C.J., and Chen, J. (1996). Different roles for CD4+ and CD8+ T lymphocytes and macrophage subsets in the control of a generalized virus infection. J Virol. 70(12), 8301–8309.

33. Karupiah, G., Coupar, B.E., Andrew, M.E., Boyle, D.B., Phillips, S.M., Mullbacher, A., Blanden, R.V., and Ramshaw, I.A. (1990). Elevated natural killer cell responses in mice infected with recombinant vaccinia virus encoding murine IL-2. J Immunol. 144(1), 290–298.

34. Karupiah, G., Fredrickson, T.N., Holmes, K.L., Khairallah, L.H., and Buller, R.M. (1993). Importance of interferons in recovery from mousepox. J Virol. 67(7), 4214–4226.

35. Karupiah, G., Woodhams, C.E., Blanden, R.V., and Ramshaw, I.A. (1991). Immunobiology of infection with recombinant vaccinia virus encoding murine IL-2. Mechanisms of rapid viral clearance in immunocompetent mice. J Immunol. 147(12), 4327–4332.

36. Kasembeli, M.M., Bharadwaj, U., Robinson, P., and Tweardy, D.J. (2018). Contribution of STAT3 to Inflammatory and Fibrotic Diseases and Prospects for its Targeting for Treatment. Int J Mol Sci. 19(8).

37. Korner, H., Cook, M., Riminton, D.S., Lemckert, F.A., Hoek, R.M., Ledermann, B., Kontgen, F., Fazekas de St Groth, B., and Sedgwick, J.D. (1997). Distinct roles for lymphotoxin-alpha and tumor necrosis factor in organogenesis and spatial organization of lymphoid tissue. Eur J Immunol. 27(10), 2600–2609.

38. Korner, H., Goodsall, A.L., Lemckert, F.A., Scallon, B.J., Ghrayeb, J., Ford, A.L., and Sedgwick, J.D. (1995). Unimpaired autoreactive T-cell traffic within the central nervous system during tumor necrosis factor receptor-mediated inhibition of experimental autoimmune encephalomyelitis. Proc Natl Acad Sci U S A. 92(24), 11066–11070.

39. Livak, K.J., and Schmittgen, T.D. (2001). Analysis of relative gene expression data using real-time quantitative PCR and the 2(-Delta Delta C(T)) Method. Methods. 25(4), 402–408.

40. Loparev, V.N., Parsons, J.M., Knight, J.C., Panus, J.F., Ray, C.A., Buller, R.M., Pickup, D.J., and Esposito, J.J. (1998). A third distinct tumor necrosis factor receptor of orthopoxviruses. Proc Natl Acad Sci U S A. 95(7), 3786–3791.

41. Ma, J.F., Sanchez, B.J., Hall, D.T., Tremblay, A.K., Di Marco, S., and Gallouzi, I.E. (2017). STAT3 promotes IFNgamma/TNFalpha-induced muscle wasting in an NF-kappaB-dependent and IL-6-independent manner. EMBO Mol Med. 9(5), 622–637.

42. Marino, M.W., Dunn, A., Grail, D., Inglese, M., Noguchi, Y., Richards, E., Jungbluth, A., Wada, H., Moore, M., Williamson, B., et al. (1997). Characterization of tumor necrosis factor-deficient mice. Proc Natl Acad Sci U S A. 94(15), 8093–8098.

43. McGeachy, M.J., Bak-Jensen, K.S., Chen, Y., Tato, C.M., Blumenschein, W., McClanahan, T., and Cua, D.J. (2007). TGF-beta and IL-6 drive the production of IL-17 and IL-10 by T cells and restrain T(H)-17 cell-mediated pathology. Nat Immunol. 8(12), 1390–1397.

44. Moss, M.L., Jin, S.L.C., Milla, M.E., Burkhart, W., Carter, H.L., Chen, W.-J., Clay, W.C., Didsbury, J.R., Hassler, D., Hoffman, C.R., et al. (1997). Cloning of a disintegrin metalloproteinase that processes precursor tumour-necrosis factor-α. Nature. 385, 733.

45. Muller, U., Steinhoff, U., Reis, L.F., Hemmi, S., Pavlovic, J., Zinkernagel, R.M., and Aguet, M. (1994). Functional role of type I and type II interferons in antiviral defense. Science. 264(5167), 1918–1921.

46. Murdaca, G., Spano, F., Contatore, M., Guastalla, A., Penza, E., Magnani, O., and Puppo, F. (2015). Infection risk associated with anti-TNF-alpha agents: a review. Expert Opin Drug Saf. 14(4), 571–582.

47. Neumann, B., Machleidt, T., Lifka, A., Pfeffer, K., Vestweber, D., Mak, T.W., Holzmann, B., and Kronke, M. (1996). Crucial role of 55-kilodalton TNF receptor in TNF-induced adhesion molecule expression and leukocyte organ infiltration. J Immunol. 156(4), 1587–1593.

48. Niemand, C., Nimmesgern, A., Haan, S., Fischer, P., Schaper, F., Rossaint, R., Heinrich, P.C., and Müller-Newen, G. (2003). Activation of STAT3 by IL-6 and IL-10 in Primary Human Macrophages Is Differentially Modulated by Suppressor of Cytokine Signaling 3. J Immunol. 170(6), 3263–3272.

49. O’Gorman, W.E., Sampath, P., Simonds, E.F., Sikorski, R., O’Malley, M., Krutzik, P.O., Chen, H., Panchanathan, V., Chaudhri, G., Karupiah, G., et al. (2010). Alternate mechanisms of initial pattern recognition drive differential immune responses to related poxviruses. Cell Host Microbe. 8(2), 174–185.

50. O’Shea, J.J., Schwartz, D.M., Villarino, A.V., Gadina, M., McInnes, I.B., and Laurence, A. (2015). The JAK-STAT pathway: impact on human disease and therapeutic intervention. Annu Rev Med. 66, 311–328.

51. Oeckinghaus, A., Hayden, M.S., and Ghosh, S. (2011). Crosstalk in NF-[kappa]B signaling pathways. Nat Immunol. 12(8), 695–708.

52. Panchanathan, V., Chaudhri, G., and Karupiah, G. (2005). Interferon function is not required for recovery from a secondary poxvirus infection. Proc Natl Acad Sci U S A. 102(36), 12921–12926.

53. Papathanasiou, S., Rickelt, S., Soriano, M.E., Schips, T.G., Maier, H.J., Davos, C.H., Varela, A., Kaklamanis, L., Mann, D.L., and Capetanaki, Y. (2015). Tumor necrosis factor-alpha confers cardioprotection through ectopic expression of keratins K8 and K18. Nat Med. 21(9), 1076–1084.

54. Parker, A.K., Parker, S., Yokoyama, W.M., Corbett, J.A., and Buller, R.M.L. (2007). Induction of natural killer cell responses by ectromelia virus controls infection. J Virol. 81(8), 4070–4079.

55. Pasparakis, M., Alexopoulou, L., Episkopou, V., and Kollias, G. (1996). Immune and inflammatory responses in TNF alpha-deficient mice: a critical requirement for TNF alpha in the formation of primary B cell follicles, follicular dendritic cell networks and germinal centers, and in the maturation of the humoral immune response. J Exp Med. 184(4), 1397–1411.

56. Pasparakis, M., Alexopoulou, L., Grell, M., Pfizenmaier, K., Bluethmann, H., and Kollias, G. (1997). Peyer’s patch organogenesis is intact yet formation of B lymphocyte follicles is defective in peripheral lymphoid organs of mice deficient for tumor necrosis factor and its 55-kDa receptor. Proc Natl Acad Sci U S A. 94(12), 6319–6323.

57. Peper, R.L., and Van Campen, H. (1995). Tumor necrosis factor as a mediator of inflammation in influenza A viral pneumonia. Microb Pathog. 19(3), 175–183.

58. Qing, Y., and Stark, G.R. (2004). Alternative activation of STAT1 and STAT3 in response to interferon-gamma. J Biol Chem. 279(40), 41679–41685.

59. Rahman, M.M., and McFadden, G. (2006). Modulation of tumor necrosis factor by microbial pathogens. PLoS Pathog. 2(2), e4.

60. Rincon, M. (2012). Interleukin-6: from an inflammatory marker to a target for inflammatory diseases. Trends Immunol. 33(11), 571–577.

61. Ruuls, S.R., Hoek, R.M., Ngo, V.N., McNeil, T., Lucian, L.A., Janatpour, M.J., Korner, H., Scheerens, H., Hessel, E.M., Cyster, J.G., et al. (2001). Membrane-bound TNF supports secondary lymphoid organ structure but is subservient to secreted TNF in driving autoimmune inflammation. Immunity. 15(4), 533–543.

62. Sanjabi, S., Zenewicz, L.A., Kamanaka, M., and Flavell, R.A. (2009). Anti-inflammatory and pro-inflammatory roles of TGF-beta, IL-10, and IL-22 in immunity and autoimmunity. Curr Opin Pharmacol. 9(4), 447–453.

63. Saunders, B.M., and Britton, W.J. (2007). Life and death in the granuloma: immunopathology of tuberculosis. Immunol Cell Biol. 85(2), 103–111.

64. Scheurich, P., Thoma, B., Ucer, U., and Pfizenmaier, K. (1987). Immunoregulatory activity of recombinant human tumor necrosis factor (TNF)-alpha: induction of TNF receptors on human T cells and TNF-alpha-mediated enhancement of T cell responses. J Immunol. 138(6), 1786–1790.

65. Schroder, K., Hertzog, P.J., Ravasi, T., and Hume, D.A. (2004). Interferon-gamma: an overview of signals, mechanisms and functions. J Leukoc Biol. 75(2), 163–189.

66. Seet, B.T., Johnston, J.B., Brunetti, C.R., Barrett, J.W., Everett, H., Cameron, C., Sypula, J., Nazarian, S.H., Lucas, A., and McFadden, G. (2003). Poxviruses and immune evasion. Annu Rev Immunol. 21, 377–423.

67. Shale, M., Czub, M., Kaplan, G.G., Panaccione, R., and Ghosh, S. (2010). Anti-tumor necrosis factor therapy and influenza: keeping it in perspective. Therap Adv Gastroenterol. 3(3), 173–177.

68. Shuai, K., and Liu, B. (2005). Regulation of gene-activation pathways by PIAS proteins in the immune system. Nat Rev Immunol. 5(8), 593–605.

69. Subramaniam, K., D’Rozario, J., and Pavli, P. (2013). Lymphoma and other lymphoproliferative disorders in inflammatory bowel disease: A review. J Gastroen Hepatol. 28(1), 24–30.

70. Szretter, K.J., Gangappa, S., Lu, X., Smith, C., Shieh, W.J., Zaki, S.R., Sambhara, S., Tumpey, T.M., and Katz, J.M. (2007). Role of host cytokine responses in the pathogenesis of avian H5N1 influenza viruses in mice. J Virol. 81(6), 2736–2744.

71. Tracey, K.J., Fong, Y., Hesse, D.G., Manogue, K.R., Lee, A.T., Kuo, G.C., Lowry, S.F., and Cerami, A. (1987). Anti-cachectin/TNF monoclonal antibodies prevent septic shock during lethal bacteraemia. Nature. 330(6149), 662–664.

72. Uysal, K.T., Wiesbrock, S.M., Marino, M.W., and Hotamisligil, G.S. (1997). Protection from obesity-induced insulin resistance in mice lacking TNF-alpha function. Nature. 389(6651), 610–614.

73. Vigano, M., Degasperi, E., Aghemo, A., Lampertico, P., and Colombo, M. (2012). Anti-TNF drugs in patients with hepatitis B or C virus infection: safety and clinical management. Expert Opin Biol Ther. 12(2), 193–207.

74. Wilhelm, P., Ritter, U., Labbow, S., Donhauser, N., Rollinghoff, M., Bogdan, C., and Korner, H. (2001). Rapidly fatal leishmaniasis in resistant C57BL/6 mice lacking TNF. J Immunol. 166(6), 4012–4019.

75. Yagil, Z., Nechushtan, H., Kay, G., Yang, C.M., Kemeny, D.M., and Razin, E. (2010). The enigma of the role of protein inhibitor of activated STAT3 (PIAS3) in the immune response. Trends Immunol. 31(5), 199–204.

76. Yamamoto, T., Matsuda, T., Muraguchi, A., Miyazono, K., and Kawabata, M. (2001). Cross-talk between IL-6 and TGF-beta signaling in hepatoma cells. FEBS Lett. 492(3), 247–253.

77. Zhong, Z., Wen, Z., and Darnell, J. (1994). Stat3: a STAT family member activated by tyrosine phosphorylation in response to epidermal growth factor and interleukin-6. Science. 264(5155), 95–98.

