## Supplemental Figures and Tables for "TNF deficiency dysregulates inflammatory cytokine production leading to lung pathology and death during respiratory poxvirus infection"

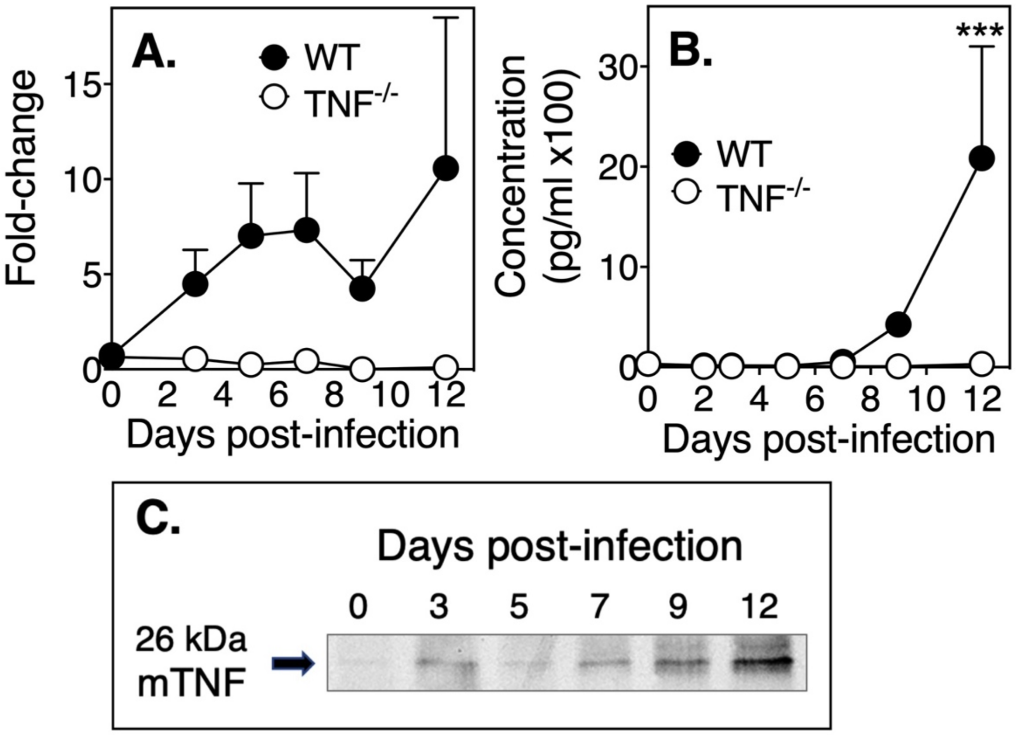


**Figure S1. Rapid induction of TNF mRNA and mTNF protein in ECTV infected WT mice.** WT and TNF$^{-/-}$ mice were infected with ECTV i.n. and on the days indicated, 4-5 mice per group were sacrificed and lung tissue was collected for RNA isolation or protein detection. (A) TNF mRNA expression was detected by real time PCR. Data are normalized to UBC gene expression from 5 mice per group and are from one experiment representative of three separate experiments. (B) Concentration of sTNF protein in lung homogenates determined by cytokine bead array from 4 mice per group. Data shown are a representative of two separate experiments. Statistics were performed using the two-way ANOVA test where ***p $<$ 0.001. (C) Western blot analysis showing mTNF (26kDa) expression by lung cells of ECTV-infected WT mice (n=8).

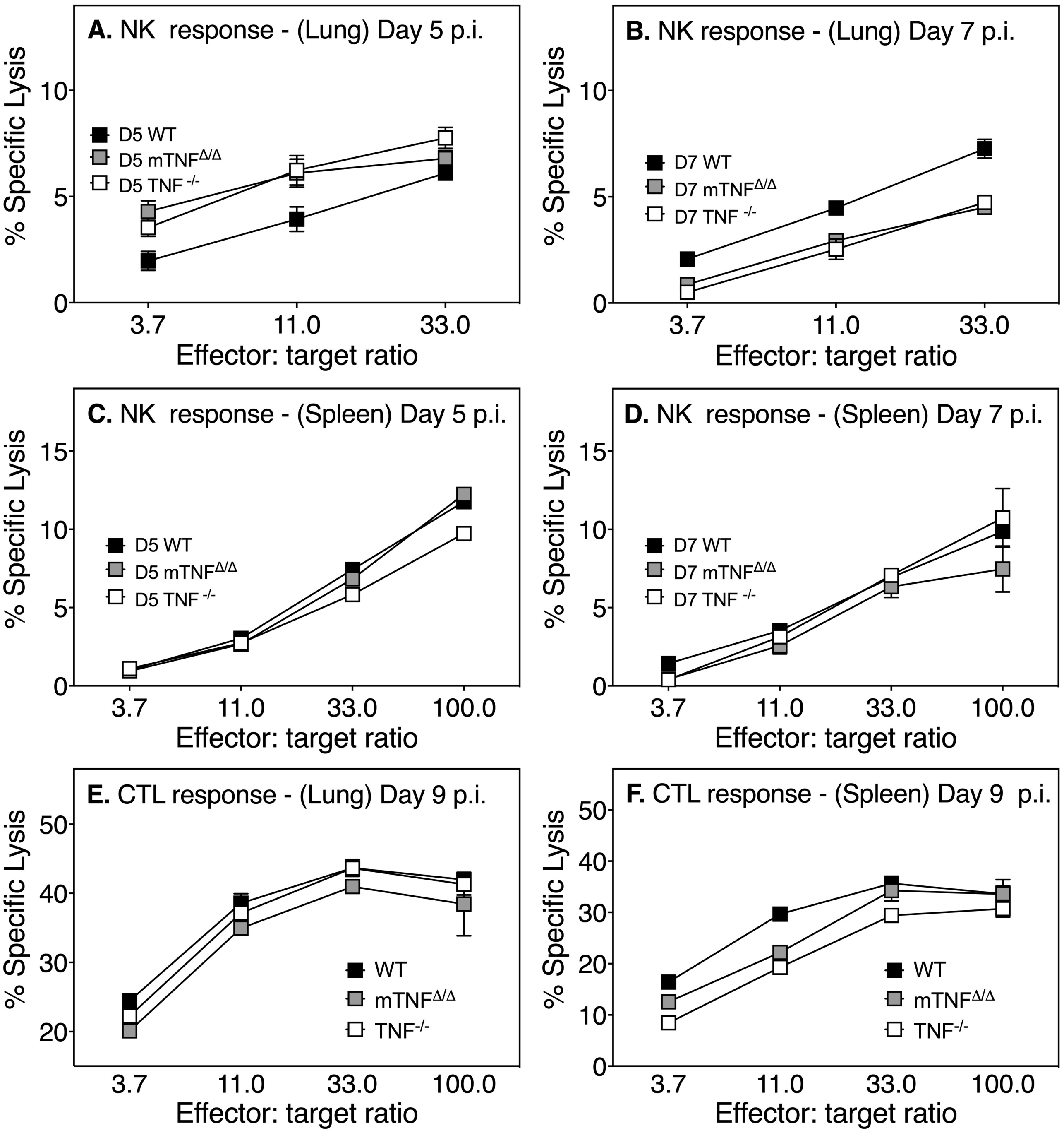


**Figure S2. NK and virus-specific cytolytic T cell activities remain intact in the absence of TNF.** The NK cell lytic activity of lung cells (A and B) and splenocytes (C and D) was measured on days 5 and 7 p.i., respectively using the NK sensitive YAC-1 target cells while the cytolytic activity of virus-specific CTL in lungs (E) and splenocytes (F) was measured on day 9 p.i. using $^{51}$Cr-labelled, ECTV-infected and uninfected MC57G target cells. Data shown is from one of two independent experiments.

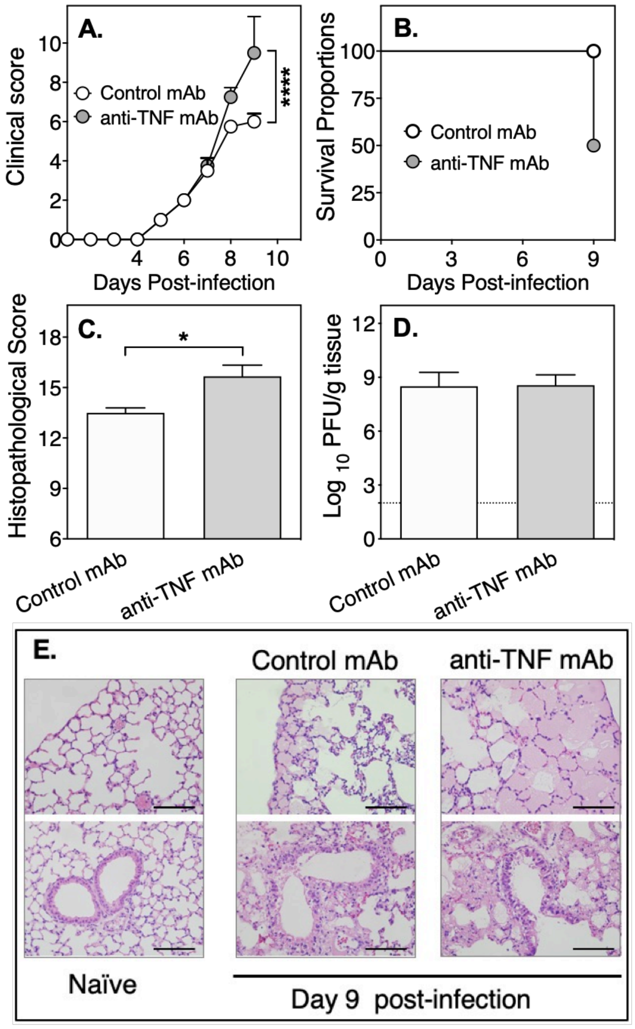


**Figure S3. Increased susceptibility of ECTV-infected C57BL/6 WT mice treated with anti-TNF mAb.** Groups of WT mice (n=4) were infected via the i.n. route with 25 PFU ECTV. At day 7 p.i., one group was treated i.p. with 500 µg anti-TNF mAb while the control group was treated with 500 µg rat IgG isotype control mAb. Two mice in the anti-TNF mAb -treated group were severely moribund at day 8 p.i. and were euthanized for ethical reasons. The remaining animals were sacrificed on day 9 p.i. and lung tissue was collected for histology and measurement of viral load. (A) Clinical scores, (B) survival proportions, (C) histopathological scores (D) viral load and (E) representative lung histology sections from naïve and infected mice. For A, significance was calculated using two-way ANOVA followed by Sidak's post-hoc tests. ****, $p < 0.0001$. For C, histopathological scores are expressed as means ± SEM and significance were determined by unpaired t-test. *, $p < 0.05$. For D, viral load data was log-transformed and the broken line indicates sensitivity of virus detection.

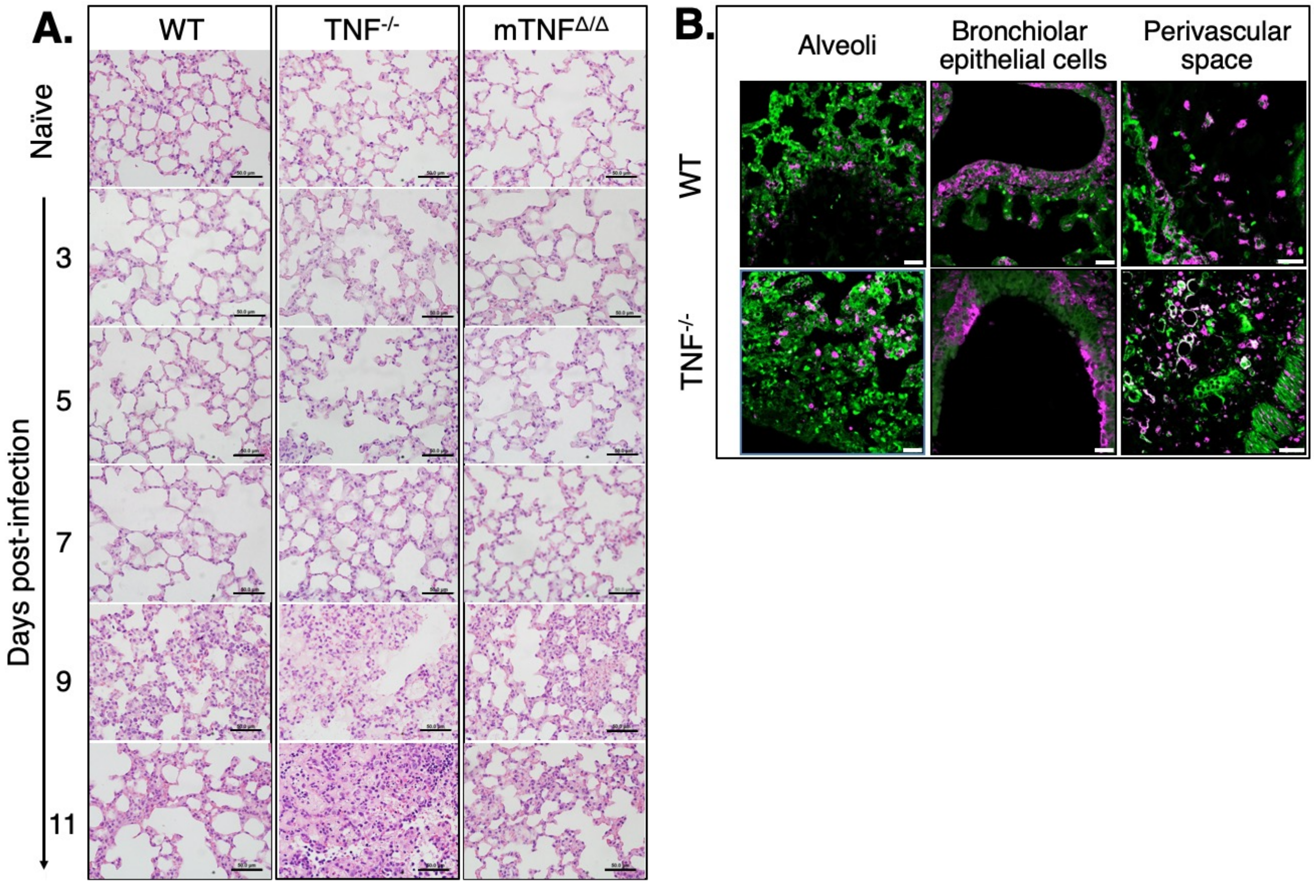


**Figure S4. Kinetics of lung pathology development in ECTV-infected WT, TNF$^{-/-}$ and mTNF$^{\Delta/\Delta}$ mice and presence of viral antigen in lungs of WT and TNF$^{-/-}$ mice.** Groups of WT, mTNF$^{\Delta/\Delta}$, and TNF$^{-/-}$ mice were infected i.n. with 25 PFU ECTV. On the days indicated, 5 animals from each strain were euthanized and lung tissue was collected for histology. (A) H&E stained lung sections from naïve and virus-infected mice sacrificed at different days p.i. Slides were examined on all fields at 400x magnification and the bars shown correspond to 100 µm. (B) Immunohistochemical staining of ECTV-infected lungs showing presence of viral antigen in WT and TNF$^{-/-}$ mice at day 9 p.i. Formalin-fixed, paraffin-embedded lung sections were stained with rabbit anti-vaccinia virus polyclonal antibody and detected using Alexa Fluor 467 conjugated to anti-rabbit IgG. Slides were examined at 600x magnification. WT and TNF$^{-/-}$ mice both exhibit similar patterns of virus spread (magenta) in alveolar spaces, bronchial epithelial cells and perivascular spaces. Auto-fluorescence from the lung tissue is shown in green. Results shown are from one of two separate experiments with similar outcomes.

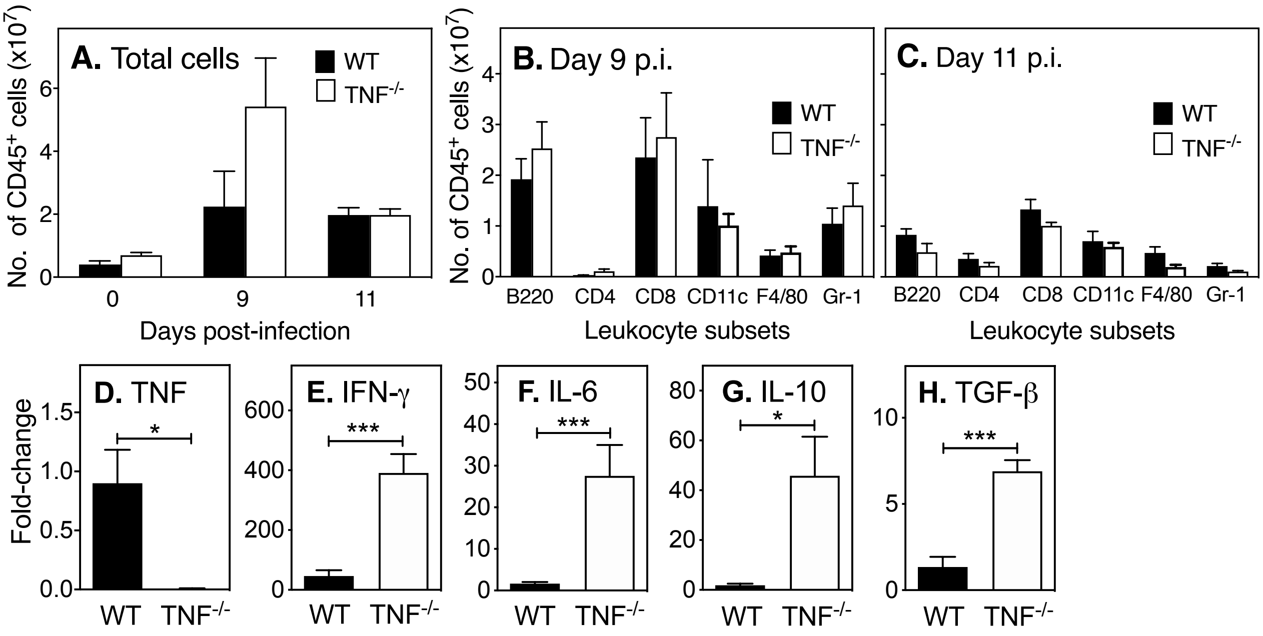


**Figure S5.** TNF deficiency does not influence lung leukocyte recruitment but results in increased expression of inflammatory cytokine mRNA transcripts. Groups of WT and TNF$^{-/-}$ mice were infected i.n. with 25 PFU ECTV and sacrificed on the days indicated. Flow cytometric analysis of digested lungs indicated that the (A) total number of leukocytes (CD45$^{+}$) or leukocyte subsets (B) and (C) were comparable between uninfected or ECTV-infected TNF$^{-/-}$ and WT strains. (D-H) Levels of expression of mRNA transcripts for IFN-γ, IL-6, IL-10, and TGF-β were significantly higher in lungs of TNF$^{-/-}$ mice compared to WT animals at day 12 p.i.

**Table S1**. Scoring system for clinical presentation of mice infected with ECTV

| | |
|---|---|
| I. | Hair Coat |
| | 0 – groomed and shiny<br>1 – groomed but not shiny<br>2 – rough<br>3 – unkempt (“scruffy”) |
| II. | Posture |
| | 0 – back is straight<br>1 – hunched but with spontaneous straightening of the back<br>2 – hunched but straightens only upon stimulation<br>3 – hunched even when stimulated |
| III. | Breathing |
| | 0 – normal<br>1 – intermittent rapid breathing<br>2 – rapid, shallow breathing<br>3 – laboured breathing; (+) abdominal retractions |
| IV. | Lacrimation and Nasal Discharge |
| | 0 – none<br>1 – minimal lacrimation or nasal discharge<br>2 – moderate lacrimation and nasal discharge<br>3 – excessive lacrimation/catarrh; (+) blockade of either nares |
| V. | Activity/Movement/Behaviour |
| | 0 – active, spontaneous movement<br>1 – inactive, movement upon stimulation<br>2 – huddled; inactive, reluctant to move<br>3 – moribund; does not go back to prone position when placed on a supine position |

**Table S2**. Histopathological assessment of lung sections from ECTV-infected animals

| | |
|---|---|
| I. | Parenchymal/Intra-alveolar Edema |
| | 0 – None |
| | 1 – Accumulation of fluid in <25% of pulmonary parenchyma |
| | 2 – Accumulation of fluid in 25-50% of pulmonary parenchyma |
| | 3 – Accumulation of fluid in 50-75% of pulmonary parenchyma |
| | 4 – Accumulation of fluid in >75% of pulmonary parenchyma |
| II. | Perivascular Edema |
| | 0 – None |
| | 1 – Mild accumulation of fluid in few perivascular spaces |
| | 2 – Mild to moderate accumulation of fluid in some perivascular spaces |
| | 3 – Moderate to severe accumulation of fluid in most of the perivascular spaces |
| | 4 – Moderate to severe accumulation of fluid in all of the perivascular spaces |
| III. | Degree of Bronchial Epithelium Necrosis |
| | 0 – None |
| | 1 – < 25% epithelial necrosis of few bronchioles |
| | 2 – < 25% epithelial necrosis of most bronchioles |
| | 3 – 25-50% epithelial necrosis of most bronchioles |
| | 4 – > 50% epithelial necrosis of most bronchioles |
| IV. | Parenchymal Inflammatory Infiltrates |
| | 0 – Few infiltrates in few areas |
| | 1 – Only few infiltrates in separate areas/foci |
| | 2 – Many scattered infiltrates in few areas/foci |
| | 3 – Confluent infiltrates in few areas/foci |
| | 4 – Diffuse inflammatory infiltrates |
| V. | Perivascular Inflammatory Infiltrates |
| | 0 – Few infiltrates in few spaces |
| | 1 – Only few infiltrates in most spaces |
| | 2 – Many scattered infiltrates in few perivascular spaces |
| | 3 – Confluent infiltrates in few perivascular spaces |
| | 4 – Infiltrates occupying most of the areas of the spaces |
| VI. | Alveolar Septal Wall Damage |
| | 0 – None |
| | 1 – Damage in <25% of the alveolar septal walls |
| | 2 – Damage in 25-50% of the alveolar septal walls |
| | 3 – Damage in 50-75% of the alveolar septal walls |
| | 4 – Damage in >75% of the alveolar septal walls |
